# *Mycoplasma gallisepticum* FtsZ demonstrates properties that distinguish it from other known homologs

**DOI:** 10.1101/2025.09.01.673419

**Authors:** Natalia A. Rumyantseva, Daria M. Golofeeva, Aizilya A. Khasanova, Anna V. Shabalina, Yana V. Bogacheva, Elena S. Ponomareva, Vasilisa S. Polynovskaya, Arina V. Drobysheva, Vadim A. Ivanov, Tatyana O. Artamonova, Alexander V. Ankudinov, Innokentii E. Vishnyakov, Mikhail A. Khodorkovskii, Alexey D. Vedyaykin

## Abstract

In bacteria, cell division usually occurs through binary fission, with the participation of genes from the *dcw* cluster. In mollicutes, this cluster is significantly reduced — often only the *ftsZ*, *ftsA*, *mraZ*, and *mraW* genes remain, and sometimes *ftsZ* is completely absent. FtsZ is a key division protein that forms the Z-ring, but its role in mollicutes is questionable due to the absence of many cell division proteins and the cell wall. In the current study, we investigated the FtsZ protein of *Mycoplasma gallisepticum*, a bacterium with a reduced set of putative cell division genes (*ftsZ*, *ftsA*, *ftsK*). The results show that, unlike in well-studied bacteria, FtsZ in *M. gallisepticum* often exhibits polar rather than mid-cell localization. Overexpression of fluorescently labeled FtsZ enhances this polar localization and may lead to minicell formation. The FtsZ concentration was measured and, together with *in vitro* data, confirmed its ability to polymerize, similar to its homologs. Protein-protein interactions were also analyzed and confirmed the link of FtsZ to cell division. Overall, the results support the role of FtsZ in cell division, though its properties differ significantly from other known homologs.

## Introduction

Cell division is a pivotal stage in the life cycle of any living organism. In bacteria, the most common mechanism is binary fission, in which the mother cell divides into two daughter cells. In their genomes, the genes encoding most proteins involved in division are concentrated in the *dcw* cluster (division and cell wall) [1]. Although this cluster is relatively conserved, the number of genes it contains can vary significantly among different bacterial species. For example, in *Escherichia coli*, the *dcw* cluster includes 15 genes, in *Bacillus subtilis* – 16, while in some mollicutes – only 4 [2–4]. Some cell division genes are typically located outside this cluster, such as *ftsK* and *ftsN*. Mollicutes lack most of the genes from the dcw cluster; for instance, *Mycoplasma pneumoniae*, *M. genitalium*, and *M. gallisepticum* retain only four genes each: *mraZ*, *mraW*, *ftsA*, and *ftsZ* [2–4]. Some mollicutes, such as *M. mobile* or *Ureaplasma spp.* [5], lack even *ftsZ* [6]. In this regard, the role of the *dcw* cluster genes involved in Z-ring formation, as well as the role of the FtsZ protein in particular, in mollicute cell division remains unclear and requires further detailed study.

FtsZ is a cytoskeletal protein present in most bacterial cells and is one of the key components of the divisome—a protein complex that mediates cell division [7]. Homologs of this protein have been found not only in bacteria but also in some archaea, as well as in plant plastids and mitochondria of certain eukaryotes [8]. Moreover, FtsZ is a homolog of eukaryotic tubulin and shares a similar 3D structure and some functional properties [9]. In well-studied bacteria, the FtsZ protein can polymerize to form single-stranded linear filaments, which may further assemble into larger structures such as bundles [10]. Polymerization of FtsZ leads to the formation of a Z-ring (or proto-ring), composed of FtsZ filaments along with anchor proteins FtsA and ZipA (in *E. coli*), which attach the filaments to the inner membrane [7]. The Z-ring subsequently recruits numerous additional proteins, forming a mature divisome that facilitates peptidoglycan synthesis, septum formation, and ultimately cell division. In other model bacteria (e.g., *B. subtilis*, Caulobacter crescentus, *Streptococcus pneumoniae*), the division mechanisms are largely similar to those in *E. coli*: FtsZ forms the Z-ring, which serves as a scaffold for other proteins involved in remodeling the cell envelope [11].

Mollicutes, however, lack a cell wall and most division proteins, retaining only FtsZ, FtsA, and a few others. This raises questions about the actual role of FtsZ in mollicutes cell division. It is hypothesized that mollicutes employ multiple division mechanisms:

1. FtsZ-dependent division (involving FtsZ and other division proteins) [12];
2. Motility-associated division (mediated by the terminal organelle) [13];
3. Membrane synthesis-driven division (due to excess membrane production) [14, 15].

Additionally, it cannot be ruled out that cells of the same species may divide in different ways, as laboratory conditions differ significantly from their natural habitats [12]. For instance, studies on *M. gallisepticum* cytokinesis have shown that these cells typically divide binary and symmetrically, forming a constriction [16]. However, cases of asymmetric polar division and variations in division site localization depending on cell shape have also been reported [17]. Division mechanisms may further depend on cell polarity. Binary fission has been observed in both polar mollicutes (e.g., *M. pneumoniae*, *M. gallisepticum*, *M. mobile* — species with a terminal organelle enabling motility) and non-polar species (e.g., *M. capricolum*). In polar mollicutes, division begins with the formation of a daughter terminal organelle, which either develops adjacent to the parent structure before migrating to the opposite pole or forms directly opposite it [17]. The presence of a terminal organelle may explain why the *ftsZ* gene is non-essential, at least in *M. pneumoniae* [13].

In this work, we investigated the properties of the FtsZ protein of *M. gallisepticum*, a bacterium with a terminal organelle and several genes encoding division proteins: *ftsZ*, *ftsA*, and *ftsK*. It was found that the FtsZ protein is localized in *M. gallisepticum* cells differently than in well-studied bacterial species: instead of predominantly localizing in the middle of the cell, FtsZ exhibits polar localization in about half of the cases. Overexpression of fluorescently labeled FtsZ promotes more pronounced polar localization of FtsZ and, presumably, the formation of minicells. The concentration of FtsZ in *M. gallisepticum* cells was determined, and, combined with data on the *in vitro* properties of this protein, it confirmed the ability of FtsZ to polymerize *in vivo*. Protein-protein interactions of the *M. gallisepticum* FtsZ protein were also characterized. The results of this work confirm the involvement of the FtsZ protein in *M. gallisepticum* cell division, although its properties significantly distinguish it from other well-studied homologs.In this work, we investigated the properties of the FtsZ protein of *M. gallisepticum*, a bacterium with a terminal organelle and several genes encoding division proteins: *ftsZ*, *ftsA*, and *ftsK*. It was found that the FtsZ protein is localized in *M. gallisepticum* cells differently than in well-studied bacterial species: instead of predominantly localizing in the middle of the cell, FtsZ exhibits polar localization in about half of the cases. Overexpression of fluorescently labeled FtsZ promotes more pronounced polar localization of FtsZ and, presumably, the formation of minicells. The concentration of FtsZ in *M. gallisepticum* cells was determined, and, combined with data on the *in vitro* properties of this protein, it confirmed the ability of FtsZ to polymerize *in vivo*. Protein-protein interactions of the *M. gallisepticum* FtsZ protein were also characterized. The results of this work confirm the involvement of the FtsZ protein in *M. gallisepticum* cell division, although its properties significantly distinguish it from other well-studied homologs.

## Materials and Methods

### Strains, Plasmids and Growth Conditions

1. *E. coli* DH5α strain was used for molecular cloning according to standard protocols (to obtain all plasmids used in the current study). The sequences of all plasmids generated in the current study are provided in the Supplementary Material (Table S1, zip archive with .gb files and Figures S1, S3–S20).

The *E. coli* Rosetta strain was used for FtsZ protein production.

For *M. gallisepticum* FtsZ production prior to purification (for antibody preparation), a plasmid based on the pBAD/HisB vector was used, into which the target gene (amplified from the genome of the corresponding species) was inserted (see Figure S1). Since the native *ftsZ* gene from *M. gallisepticum* contains two non-canonical tryptophan codons (TGA, which are stop codons in *E. coli*), site-directed mutagenesis was performed via PCR to enable full-length gene expression. This resulted in nucleotide substitutions (TGA→TGG) in the codons corresponding to amino acids W322 and W416. The resulting plasmid map is shown in Figure S1. Overnight cultures were diluted 1:100 in fresh SOB medium containing 100 μg/ml ampicillin. When OD600 reached ∼0.5, L-arabinose was added to a final concentration of 10 mM. Induction was carried out for 12 h at 18°C.

For *M. gallisepticum* FtsZ production prior to purification (for *in vitro experiments*, including polymerization and GTPase activity assays), cells were transformed with an expression vector based on the pET21a plasmid containing the *ftsZ* gene with optimized codons (see Figure S3). Codon composition was optimized using the IDT DNA codon optimization tool. The *ftsZ* gene was synthesized *de novo* by IDT DNA. Overnight cultures were diluted 1:100 in fresh SOB medium containing 100 μg/ml ampicillin. When OD600 reached ∼0.5, IPTG was added to a final concentration of 1 mM. Induction was carried out for 12 h at 18°C.

An *E. coli* Δ*cyaA* mutant derived from the K802 strain (custom-made in the laboratory) was used for the bacterial two-hybrid assay. To generate this strain, the wild-type K802 strain was transduced with P1 phage previously propagated on the Δ*cyaA* strain from the Keio collection. Plasmids were obtained using standard techniques. The *M. gallisepticum* genes were mutated (TGA→TGG) as described for *ftsZ*. The sequences of all plasmids used in the two-hybrid assay are shown in Figures S3–S20. To enhance *M. gallisepticum* protein expression in *E. coli*, a plasmid encoding tRNAs corresponding to rare codons (see Figure S20) was used for bacterial transformation.

The *M. gallisepticum* S6 strain was used for microscopy and co-immunoprecipitation in this work. All manipulations were performed as described previously [18–20]. Briefly, the bacteria were cultivated for 24 h at 37°C without shaking until the exponential growth phase was reached. The cultures were grown in a liquid medium consisting either of Mycoplasma broth base (OXOID, GB) supplemented with G (OXOID, GB) or a custom medium (2% tryptose, 0.5% NaCl, 0.3% Tris, 0.13% KCl, 5% homemade yeast dialysate, 10% horse serum) supplemented with sterile glucose (final concentration 1%) and phenol red (0.04%). The transition to the exponential phase was determined by a color change of the indicator from red to yellow due to medium acidification.

For fluorescent endogenous labeling of the FtsZ protein, a plasmid based on the pTn4001mini_Recipient-MCS-M5 vector was constructed (see Figure S4). This vector contains a transposase gene, an ampicillin resistance gene, a transposon carrying a tetracycline resistance gene, and a multiple cloning site. The gene of interest and the mMaple2 fluorescent protein gene were inserted into the multiple cloning site. mMaple2 was chosen because it is photoswitchable and suitable for SMLM microscopy. Upon excitation with 405 nm light, mMaple2 shifts from a green to a red fluorescent state.

For transformation, 10 mL of an exponential-phase *M. gallisepticum* S6 culture was harvested. The cells were pelleted by centrifugation (10,000 × g, 10 min, 0°C). The supernatant was carefully removed, and the cells were washed twice with 10 mL of ice-cold buffer (8 mM HEPES, 272 mM sucrose, pH 7.4). The cells were then transferred to a 1.5 mL tube, centrifuged again (10,000 × g, 10 min, 0°C), and resuspended in 250 μL of buffer. Transformation was performed via electroporation. 3–5 μg of plasmid DNA was added to 50 μL of cell suspension, and the mixture was transferred to a pre-chilled electroporation cuvette. Electroporation was carried out using a MicroPulser electroporator (Bio-Rad) at 2.5 kV. The cells were recovered in 1 mL of mollicutes growth medium and incubated for 2 h at 37°C. To select transformed clones, the cells were plated in semi-solid medium. For preparation, 10 mL of mollicutes medium was heated to 42°C in a water bath. 1 mL of melted 4% agar (cooled to 42°C) was added to the medium along with tetracycline (2 mg/mL) as a selective marker. 1 mL of transformed cells was mixed into the medium, and the mixture was left in the dark at room temperature for 30 min to solidify. The tubes were then incubated at 37°C for 1–2 weeks. Colonies from the semi-solid medium were transferred to 1 mL of liquid mollicutes medium containing tetracycline and cultured at 37°C for 2–4 days. The cultures were then subcultured in fresh medium. Exponential-phase cells were either used for fluorescence visualization or frozen in liquid nitrogen and stored at –80°C.

### Purification of FtsZ Protein for Antibody Preparation

Strep-Tag purification of *M. gallisepticum* FtsZ was performed as follows. Before application, protein fractions were transferred to buffer W1 (100 mM Tris/HCl, pH 8.0, 150 mM NaCl, 1 mM EDTA) and then loaded onto a 1 ml Strep-Tactin Superflow HC column (IBA) at a flow rate of 0.5 ml/min (the same flow rate was maintained in subsequent steps). The column was washed with buffer W1, and elution was carried out using buffer E1 (100 mM Tris/HCl, pH 8.0, 150 mM NaCl, 1 mM EDTA, 2.5 mM D-desthiobiotin). The purified protein was analyzed by SDS-PAGE and concentrated by diafiltration. FtsZ was stored at −20°C in the presence of 50% glycerol.

### Preparation of Custom Antibodies

Polyclonal antibodies against the purified recombinant full-length FtsZ protein from *M. gallisepticum* were generated as described in [21]. Serum containing polyclonal antibodies against *M. gallisepticum* FtsZ was enriched by passing it through a CNBr-activated Sepharose 4B column (GE Healthcare) with immobilized target protein, following a standard protocol. Immunoglobulins used as a negative control were isolated from non-immunized rabbit serum using Protein A Sepharose (GE Healthcare) according to the manufacturer’s instructions. All custom antibodies were validated by immunoblotting.

### Quantification of FtsZ in *M. gallisepticum* Cells

The concentration of FtsZ in *M. gallisepticum* cells was estimated by semi-quantitative Western blotting. The cell count in the medium was determined using light microscopy by counting cells in a fixed volume of suspension. A lysate from a known number of cells was loaded into one lane of the gel, while increasing amounts of purified recombinant FtsZ (0.1–100 ng) were loaded into adjacent lanes.

Primary antibodies (polyclonal anti-*M. gallisepticum* FtsZ, generated in this study) were used at a 1:1000 dilution, and secondary antibodies (horseradish peroxidase-conjugated anti-rabbit immunoglobulins) were used at 1:20,000. Chemiluminescence was detected using the SuperSignal West Pico Chemiluminescent Substrate kit. The total chemiluminescence intensity of the band corresponding to the target protein was measured, ensuring that imaging conditions prevented signal saturation in the gel documentation system. Cell volume was estimated based on electron microscopy images.

### Immunofluorescence Microscopy

*M. gallisepticum* cells of the S6 strain were cultured as described above until the middle of the exponential growth phase. Cell fixation was performed in the medium by adding formaldehyde to a final concentration of 2% and glutaraldehyde to 0.1% for 1 h at room temperature. The cells were then washed once with PBS and subsequently fixed in wash chambers prepared as described in [22]. The chambers, filled with the mycoplasma cell suspension, were centrifuged at 1000 × g for 5 min, resulting in cell sedimentation and attachment to the cover glass surface. For permeabilization, a 0.1% solution of Triton X-100 in PBS was added to the chamber and incubated for 5 min. To block nonspecific antibody binding, the cells were incubated in a 3% solution of skim milk powder in PBS supplemented with 5% horse serum (to block immunoglobulin-binding proteins) for 1 h at room temperature. Primary antibodies, prepared as described above, were added at a 1:100 dilution in a non-fat dry milk solution and incubated overnight at 4°C. Washing, secondary antibody incubation, and DNA staining were performed as follows:

- The cells were washed five times with 50 µL of a 0.01% Tween-20 solution in PBS.
- Incubation with secondary antibodies conjugated to Alexa Fluor 647 (Life Technologies) at a 1:500 dilution was carried out for 1 h at room temperature, followed by washing as described above.
- For enhanced fluorescence stability, samples were post-fixed with 2% formaldehyde in PBS for 20 min at room temperature.
- For DNA visualization, a 100 nM YOYO-1 solution in PBS was added for 10 min.

- For membrane staining, cells were incubated with CellMask Green dye (Life Technologies) at concentrations ranging from 100 to 1000 nM in PBS for 10 min.

The solution proposed in [23] was added to the wash chambers containing M. gallisepticum cells. This solution was based on PBS-Tris buffer (10 mM, pH 7.5) supplemented with: 10% glucose, 10 mM β-mercaptoethylamine, 50 mM β-mercaptoethanol, 2 mM cyclooctatetraene, 2.5 mM protocatechuic acid (Sigma-Aldrich 37580), 50 nM protocatechuate dioxygenase (Sigma-Aldrich P8279). The samples were then sealed with an appropriate sealant and placed in a microscope equipped for localization microscopy, as previously described [24, 22]. For Alexa Fluor 647 excitation in localization microscopy mode, a 640 nm diode laser was used, generating a spot with a power density of ∼1 kW/cm² in the sample plane. In some cases, to accelerate Alexa Fluor 647 transitions from the dark to the fluorescent state, additional illumination with a 405 nm diode laser at low power density (1–100 W/cm²) was applied. For standard (diffraction-limited) microscopy, low-intensity light from a mercury lamp was used. The following filter sets were employed for fluorescence separation:

- Filter set 10 (Carl Zeiss) for CellMask Green and YOYO-1;
- LF635/LP-B-000 (Semrock) for Alexa Fluor 647.

In localization microscopy mode, typically 1000–20,000 consecutive frames of single-molecule fluorescence were captured and processed using the ThunderSTORM plugin [25] for ImageJ [26]. The following reconstruction parameters were applied: Intensity > 1000, Uncertainty < 10.

In conventional (diffraction-limited) fluorescence and transmitted light microscopy, 10–20 consecutive images were acquired and averaged to improve contrast. Background subtraction was performed in some cases.

### Fluorescence Microscopy of Living *M. gallisepticum* Cells

*M. gallisepticum* cells expressing the FtsZ fusion protein with the mMaple2 fluorescent protein were seeded in fresh nutrient medium at a 1:5 ratio and cultured for 1 h at 37 °C without shaking. To visualize DNA, Hoechst 33342 dye was added at a concentration of 100 nM, followed by incubation for an additional 30 min at 37 °C. The cells were then transferred to a microculture chamber formed by a coverslip and a slide separated by double-sided tape. To improve mollicute fixation, the coverslips were pre-coated with poly-L-lysine. Imaging was performed automatically at 30-min intervals using a thermostated microscope (Nikon Ti-E).

### Immunoelectron Microscopy

Cell culturing and fixation were performed similarly to the preparation for immunofluorescence microscopy. After fixation, the cells were collected by centrifugation (10,000 × *g* for 10 min at 4 °C). The pellets were dehydrated in increasing concentrations of ethanol (70% for 15 min, 96% for 15 min) and then sequentially impregnated with LR-White acrylic resin (Polysciences) in ethanol at ratios of 1:3 (2 h), 1:1 (2 h), and 3:1 (2 h). After an additional 2 h in pure LR-White resin, the pellets were transferred to gelatin capsules for embedding. Resin polymerization was carried out at 50–52 °C for 2 days. Ultrathin sections were prepared using an LKB-III ultratome (LKB, Sweden).

The sections were placed on collodion-coated nickel grids and treated with anti-FtsZ antibodies (1:100 dilution in PBS containing 0.1% BSA). Instead of secondary antibodies, a protein A conjugate with 10 nm colloidal gold particles (EY Laboratories, USA) was used. The sections were contrasted with Uranyl Acetate Alternative (Ted Pella, USA) for 10–20 min at room temperature. Electron microscopy was performed on a Libra 120 microscope at the Institute of Cytology of the Russian Academy of Sciences, following the protocol described in reference [27].

### Co-Immunoprecipitation (Co-IP) Analysis

Co-IP was performed using protein A-Sepharose resin (GE Healthcare) and affinity-purified anti-FtsZ antibodies, as previously described [28], with minor modifications. For Co-IP, protein A-Sepharose (GE Healthcare) was cross-linked with two types of antibodies: custom-made rabbit polyclonal antibodies against *M. gallisepticum* FtsZ and normal rabbit serum as a negative control. The normal serum served as a negative control since it was derived from a non-immunized rabbit and is commonly used for such purposes, including immunocytochemistry [29]. Cross-linking was performed as described previously [28]. The Co-IP procedure was as follows: *M. gallisepticum* FtsZ-expressing cells were washed twice in PBS containing 0.3 M sucrose and lysed in PBS with 1 mg/mL lysozyme (Sigma-Aldrich, UK), supplemented with 1 mM PMSF. The cells were disrupted by sonication, and the lysate was centrifuged (15,000 × g, 10 min, 4 °C) to remove debris and unbroken cells. The supernatant was diluted with NET buffer I (50 mM Tris-HCl (pH 7.0), 150 mM NaCl, 0.1% (v/v) Nonidet P-40, 1 mM EDTA) and filtered through a 0.22 μm filter. The clarified lysate was then incubated with antibody-coupled protein A-Sepharose for 4 h at 4 °C. After incubation, the Sepharose beads were washed three times with NET buffer I, twice with NET buffer II (NET buffer I containing 500 mM NaCl), and once with buffer IP (10 mM Tris-HCl (pH 7.5), 0.1% (v/v) Nonidet P-40). Bound proteins were eluted with buffer IE (100 mM glycine, pH 2.4), concentrated, and analyzed by SDS-PAGE followed by mass spectrometry [30].

### Purification of FtsZ Protein for *In Vitro* Assays

The FtsZ protein from *M. gallisepticum* was purified using a protocol similar to the standard procedure for isolating *E. coli* FtsZ protein [31]. Cells were harvested and suspended in cold lysis buffer (50 mM Tris, pH 8.0; 100 mM NaCl; 1 mM EDTA; 1 mM PMSF). Lysozyme was added to the cell suspension at a concentration of 1 mg/ml, and the cells were lysed using an ultrasonic homogenizer. Cell debris was removed by centrifugation at 30,000 × g for 30 min.

For preliminary purification of the FtsZ protein, the ammonium sulfate precipitation method was used. The supernatant was treated with 10% ammonium sulfate, incubated on ice for 20 min, and the protein was pelleted by centrifugation at 30,000 × g at 4 °C for 10 min. The pellet was resuspended in TKEM buffer (50 mM Tris, pH 8.0; 100 mM KCl; 1 mM EDTA; 5 mM MgCl). The ammonium sulfate concentration was then increased to 20%, and the steps were repeated. Two subsequent purification steps involved reprecipitation with ammonium sulfate followed by centrifugation at 100,000 × g for 20 min to remove insoluble contaminants.

The resulting protein fractions were further purified by gel filtration (Superdex® 75 10/300 GL). The column was pre-equilibrated with TKEM buffer, and chromatography was performed using the Äkta Pure 10 system. Purified protein fractions were concentrated using Amicon® filters. Total protein concentration was determined by the Bradford method (Bio-Rad Protein Assay). Proteins were flash-frozen in liquid nitrogen and stored at −80 °C.

### Analysis of FtsZ Protein Polymerization by Light Scattering

Protein polymerization was monitored by 90° light scattering. Acrylic cuvettes containing a mixture of protein and polymerization buffer were placed in a Varian Cary Eclipse spectrofluorimeter (settings: excitation wavelength, 350 nm; emission wavelength, 350 nm; excitation slit, 5 nm; emission slit, 5 nm; detector voltage, low) at 30 °C. After confirming a stable scattering signal, GTP was added to a final concentration of 3 mM.

Polymerization was assayed in the following buffers:

1. 50 mM MES (pH 6.5), 50 mM K, 5 mM MgCl, 1 mM EDTA;
2. 50 mM MOPS (pH 7.0), 50 mM K, 5 mM MgCl, 1 mM EDTA;
3. 50 mM Tris-HCl (pH 7.5), 50 mM K, 5 mM MgCl, 1 mM EDTA;

For MES- and MOPS-based buffers, pH was adjusted with KOH, and the K concentration was brought to 50 mM by adding KCl. NaOH was avoided because Na ions can destabilize FtsZ protein dimers [32]. To optimize polymerization conditions, Mg² and K concentrations in the MES-based buffer (1) were varied by supplementing with MgCl and KCl, respectively.

### Visualization of FtsZ Protein Polymerization by AFM

FtsZ polymers adsorbed onto freshly cleaved mica were visualized using an Integra Aura atomic force microscope (AFM) (NT-MDT, Zelenograd, Moscow). A 50-μL sample containing MOPS buffer (50 mM MOPS, pH 7.0; 50 mM K (KCl + KOH); 5 mM MgCl; 1 mM EDTA), FtsZ (2.5 μM), and GTP (5 mM) was applied to the mica surface, incubated for 5 min, and washed with GTP-free buffer.

Measurements were performed in a liquid cell using hybrid AFM mode (analogous to PeakForce QNM) with CSG10 cantilevers (spring constant, ∼0.3 N/m; tip radius, ∼10 nm). Scanning parameters were as follows: Probing frequency and amplitude: 250 Hz, 50 nm; Peak force: 0.2 nN; Line scan frequency: 0.25 Hz.

### Visualization of FtsZ Protein Polymerization by EM

Protein samples in the appropriate buffer (the same as those analyzed by light scattering) were applied to collodion-coated nickel grids that had been pre-treated with ultraviolet radiation for 1 min to remove electrostatic charge. After 15 sec, the buffer was removed using filter paper, and Uranyl Acetate Alternative was applied to the grid. Following another 15 sec, the contrast agent was blotted away with filter paper, and the grids were briefly rinsed with distilled water before being dried. The processed grids were examined using a Libra 120 electron microscope (Carl Zeiss, Germany) at a magnification of 10,000–20,000.

### GTPase activity assay

To measure GTPase activity, the FtsZ protein (final concentration: 5 μM) was mixed with the reaction buffer (containing 1 mM GTP) and incubated at 30 °C. At specific time points (0, 10, 20, and 30 min), 200 μl of malachite green solution (final concentration: 40 μl) was added, followed by incubation at room temperature (22 °C) for 30 min. Notably, in replicate experiments, the volumes and concentrations of reagents remained unchanged, with the key variable being the incubation time of the protein with GTP prior to inhibitor addition. The optical density of the solutions was measured at 630 nm using a platereader (BMG ClarioStar). GTP hydrolysis curves were plotted based on the obtained data, and the GTPase activity of the protein fractions was calculated (± standard deviation).

### Bacterial Two-hybrid Assay

Protein-protein interactions were analyzed using the chromogenic substrate o-nitrophenyl-β-D-galactopyranoside (ONPG). Bacterial cultures were grown to an OD600 of 0.3–0.4, after which 0.5 mM IPTG was added to induce protein expression. Following a 1-hour incubation at 37 °C, cells were lysed by adding 30 μl of toluene and 30 μl of SDS. The enzymatic reaction with ONPG was performed in PM2 buffer (70 mM Na HPO ·12H O, 30 mM NaH PO ·H O, 1 mM MgSO, 0.2 mM MnSO) until a yellow color developed, at which point the reaction was quenched with Na CO. The OD420 of each sample was measured using a plate reader (BMG ClarioStar), and β-galactosidase activity was calculated. Activity (in conventional units per milligram of dry bacterial mass) was determined using the formula:

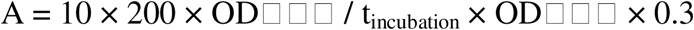

## Results and Discussion

### 1. Visualization of FtsZ in *M. gallisepticum* Cells

Immunofluorescence labeling using custom antibodies against the *M. gallisepticum* FtsZ protein successfully visualized the localization of this protein in fixed cells (see Figure 1). The results indicate that, in most cells during the exponential growth phase, FtsZ is not localized at the midcell (unlike well-studied bacterial species such as *E. coli* and *B. subtilis* [33, 34]). Structures resembling Z-rings were observed only in a small fraction of cells, whereas in most cells, FtsZ exhibited either uniform or polar localization. Notably, even in cases of uniform distribution, FtsZ formed distinct patches or clusters, similar to those observed in other bacteria [22]. The absence of Z-rings in most cells may suggest either a distinct mechanism of FtsZ involvement in cell division in *M. gallisepticum* (if FtsZ indeed participates in this process), or that only a small fraction of cells are actively dividing at any given time. Further clarification requires dynamic visualization in living cells (see below).

**Figure 1.**
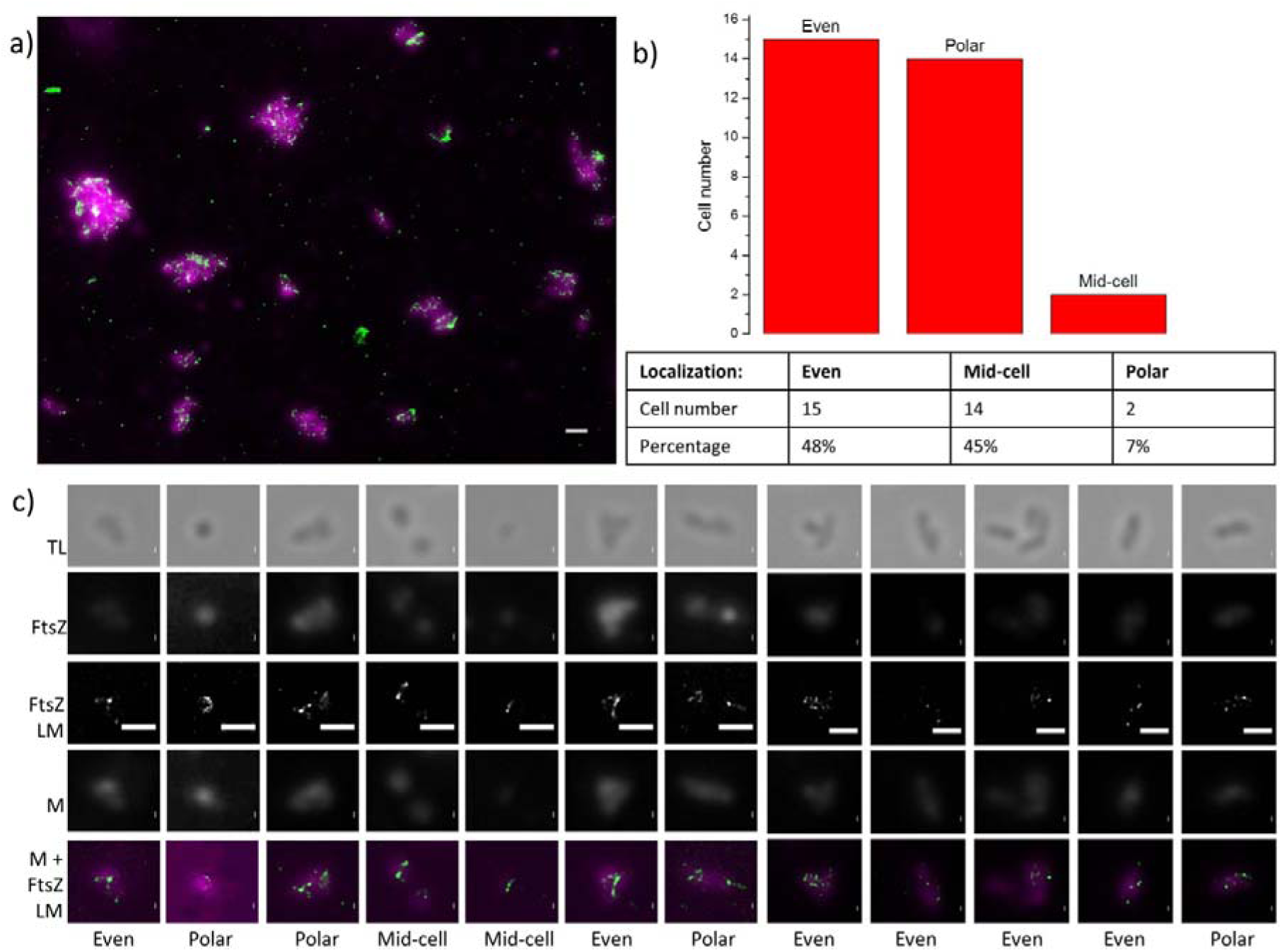
Visualization of fixed *M. gallisepticum* cells using immunofluorescence microscopy. a) Field of view of fixed cells attached to a coverslip, showing FtsZ (SMLM microscopy, green) and the membrane (standard fluorescence microscopy, magenta). b) Distribution of FtsZ localization patterns in a sample of 31 cells: even, polar, and mid-cell. c) Individual *M. gallisepticum* cells exhibiting different FtsZ localization patterns (even, polar, and mid-cell), visualized in: Transmitted light (TL), Fluorescent channel of anti-FtsZ antibodies (FtsZ – conventional fluorescence microscopy; FtsZ LM – SMLM microscopy), Fluorescent channel of the membrane dye (M), Combined channel of FtsZ LM (green) and M (magenta). Scale bar: 1 μm.

Similar results were obtained by electron microscopy (see Figure 2). The visualization data indicate that FtsZ behavior in *M. gallisepticum* differs significantly from that in bacteria exhibiting “classical” division with Z-ring formation. FtsZ in mycoplasma cells likely does not assemble into a typical Z-ring. The figure illustrates representative FtsZ localization patterns, though it does not reflect their relative frequencies. Only in a subset of *M. gallisepticum* cells was FtsZ localized at the midcell, whereas the vast majority exhibited either uniform distribution of immunogold particles or their accumulation at one of the cell poles.

**Figure 2.**
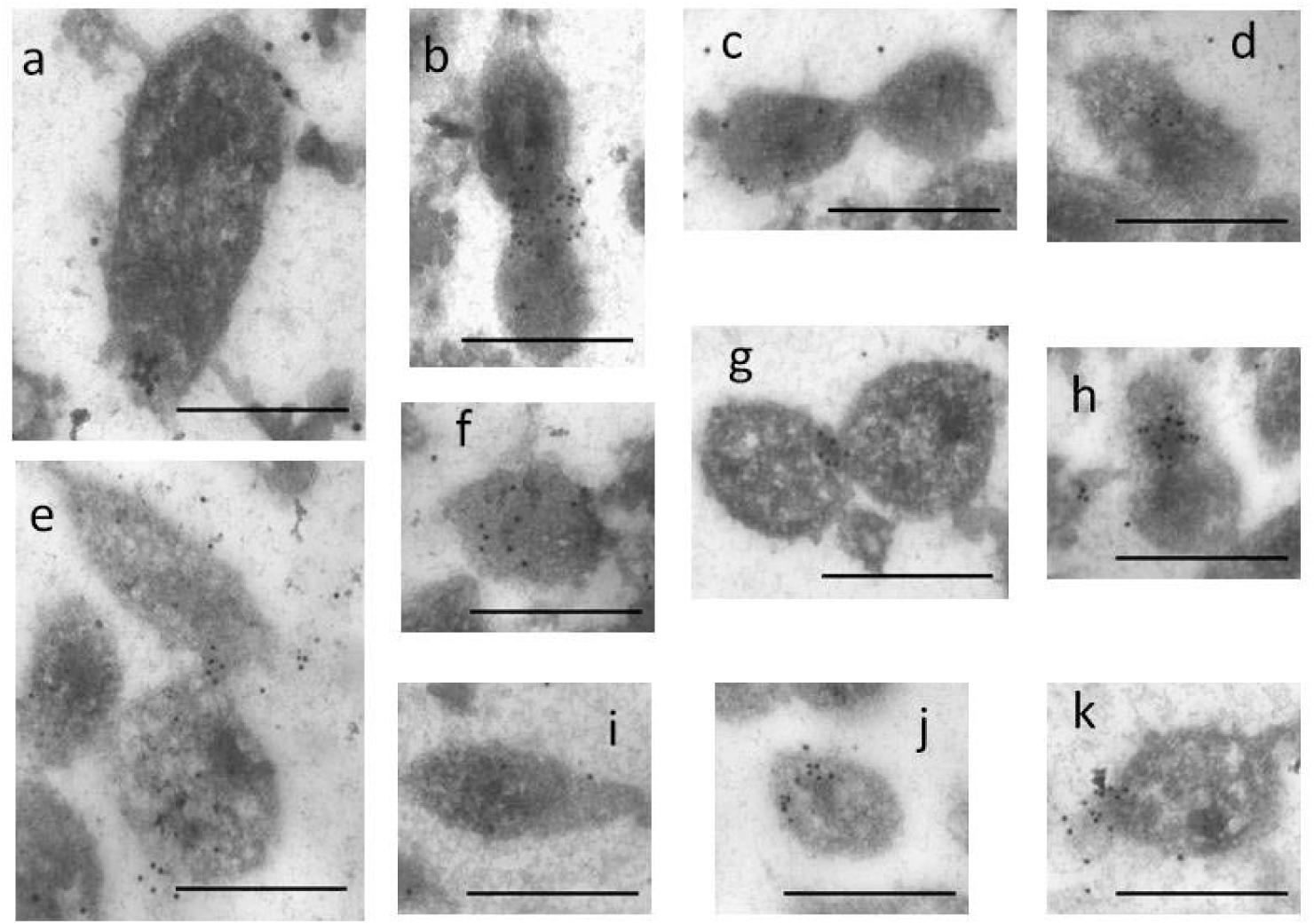
Visualization of fixed *M. gallisepticum* cells using immunoelectron microscopy. In images (a, h, j, k), gold particles show polar localization; in (b, d, e, g), mid-cell localization; and in (c, f, i), nearly uniform distribution. Note that these images do not represent the frequency of these patterns. Scale bar: 500 nm

Thus, the results of immunofluorescence and immune-electron microscopy indicate that FtsZ in *M. gallisepticum* cells is localized differently than in the “classical” case (*E. coli*, etc.). However, given the limitations of methods based on the use of antibodies and chemically fixed cells, we performed visualization using endogenous labeling to further verify the obtained results. The genetic construct integrated into the *M. gallisepticum* genome contained the FtsZ fusion gene with the mMaple2 fluorescent protein, which enabled the production of fluorescently labeled FtsZ. As a result, the construct encoding fluorescently labeled FtsZ was assembled and cloned into *M. gallisepticum* cells, allowing us to visualize the distribution of the labeled FtsZ in mollicutes cells *in vivo* for the first time.

Analysis of the visualization results revealed an uneven distribution of FtsZ in *M. gallisepticum* cells. Specifically, in most cells, FtsZ was localized at one of the poles (see Figure 3), which contrasts with the antibody-based visualization data. This discrepancy might be due to the high expression level of the fluorescently labeled FtsZ (where the protein amount was several times greater than that of the native protein — data not shown) and represents a significant limitation of the approach. However, despite this limitation, the cells exhibited a normal growth rate and morphology that was virtually indistinguishable from the wild type, prompting us to proceed with this approach for further studies. Moreover, we developed a genetic construct with a fusion protein gene under the control of a native promoter, which provided expression of the fluorescently labeled protein at a level comparable to the native one. However, this construct could not be used for dynamic visualization due to an insufficient fluorescence signal.

**Figure 3.**
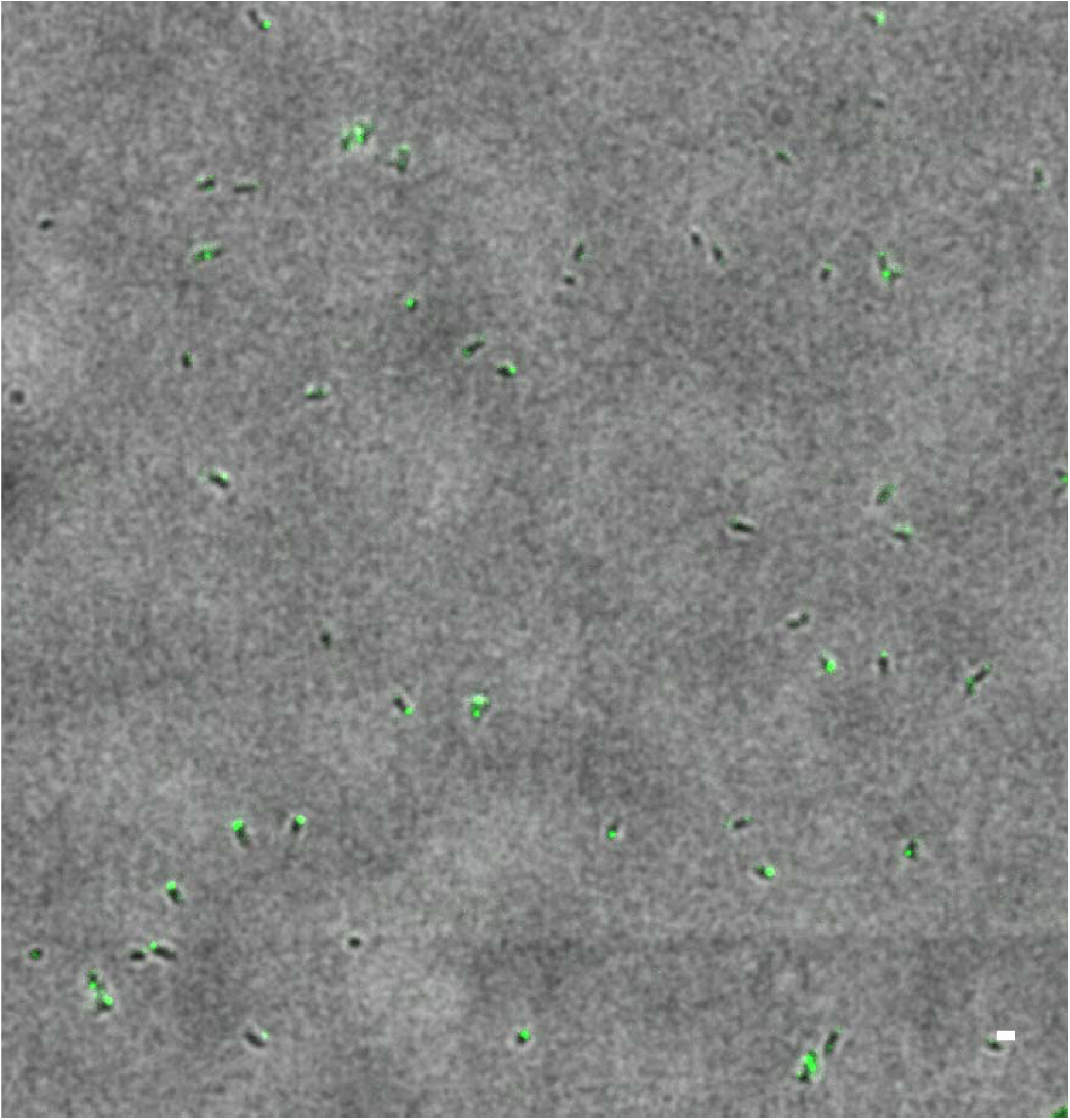
Images of structures formed by the FtsZ protein in living *M. gallisepticum* cells, obtained using the fluorescent protein mMaple2 (green). To visualize cell morphology, the diffraction-limited fluorescent image of FtsZ is combined with a transmitted light image (grayscale). Scale bar: 1 μm

Dynamic imaging of FtsZ in living *M. gallisepticum* cells revealed that, at different stages of cell division, FtsZ localizes both at the pole and at the constriction between future daughter cells (see Figure 4). Furthermore, in some cases, simultaneous imaging of FtsZ and DNA revealed the formation of DNA-free “mini-cells” containing large amounts of FtsZ (see Figure 5). The appearance of such mini-cells is likely due to the overproduction of FtsZ, which promotes cell detachment, possibly through a budding-like mechanism.

**Figure 4.**
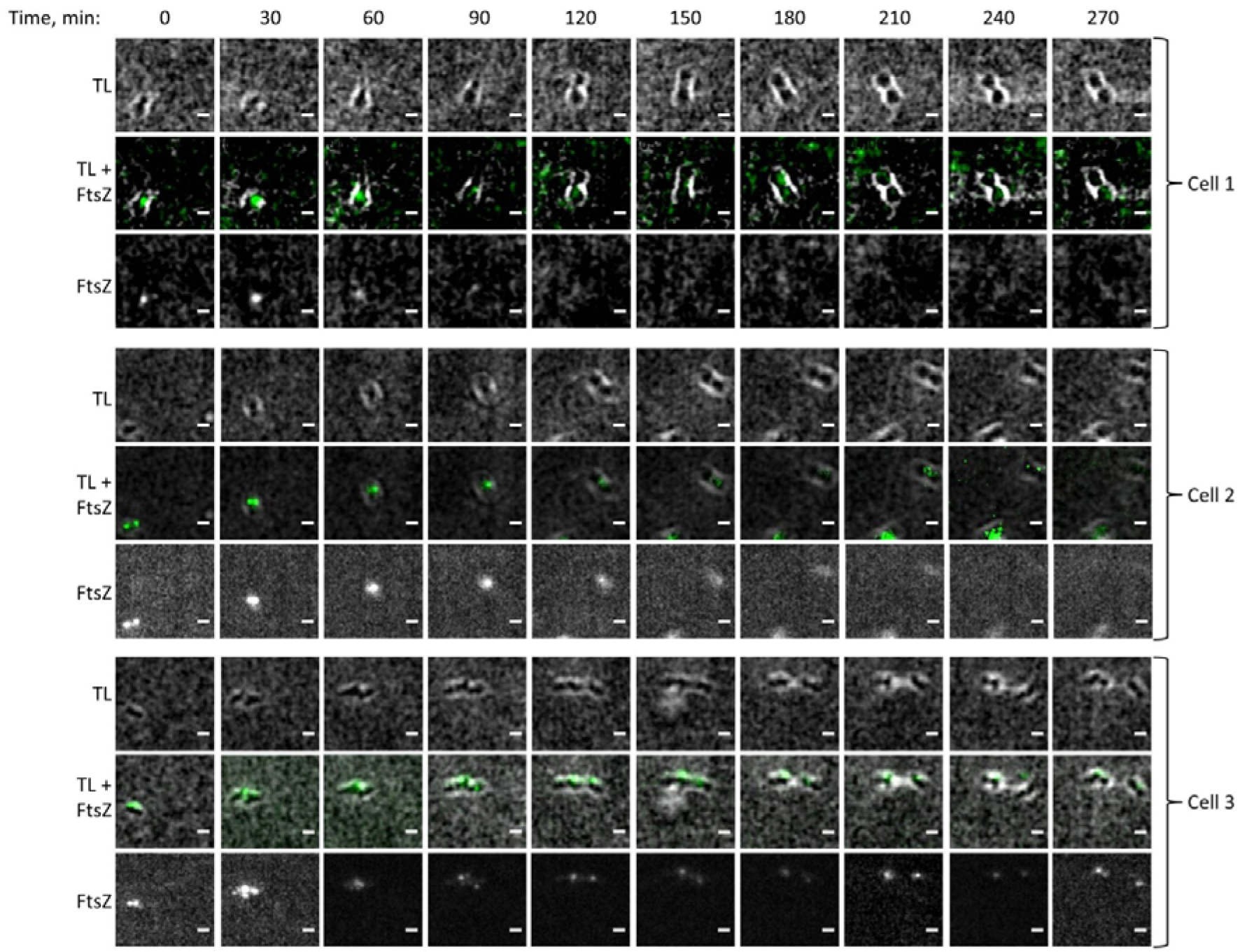
Visualization of living *M. gallisepticum* cells using time-lapse fluorescence microscopy. Sequential images at 30-minute intervals for three individual cells are shown in transmitted light (TL), FtsZ:mMaple2 fluorescence (FtsZ) channels, and a combined TL + FtsZ image. Scale bar: 1 μm

**Figure 5.**
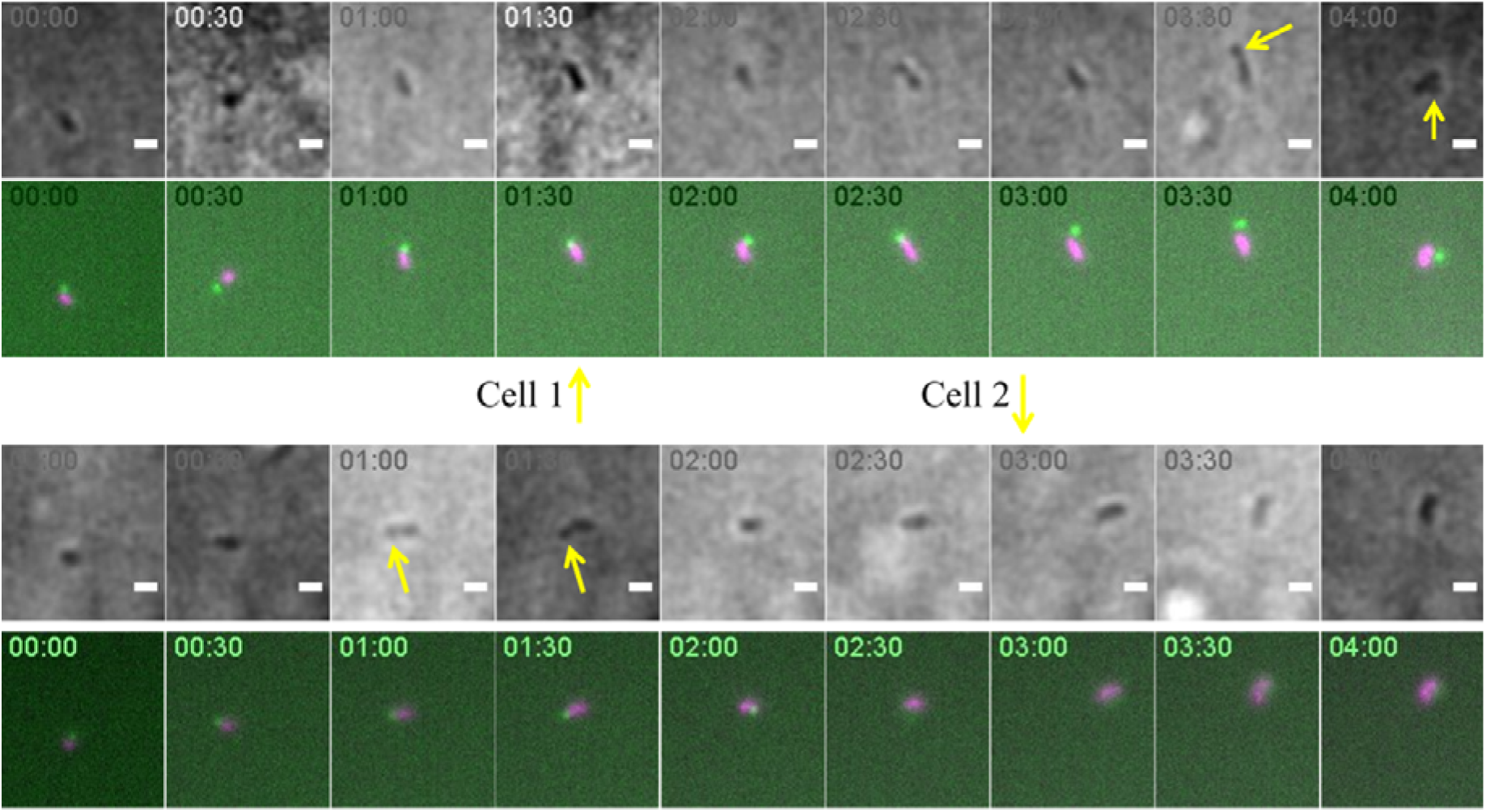
*M. gallisepticum* cells producing the FtsZ::mMaple2 fusion (overexpressed at several times the WT FtsZ concentration) demonstrate nearly normal cell division and growth rates. FtsZ is shown in green, DNA in magenta. Unlike FtsZ in WT cells visualized by immunofluorescence (IF), FtsZ localizes at the poles of most cells. Arrows indicate probable “mini-cells” that lack DNA but contain abundant FtsZ. Their formation is presumably due to a budding-like mechanism facilitated by FtsZ. Scale bar: 1 μm

These observations suggest that FtsZ is involved in *M. gallisepticum* cell division, though its role differs from the classical divisome-associated function: instead of forming a contractile ring, FtsZ appears to accumulate at one of the cell poles and facilitates daughter cell detachment.

### 2. Estimation of FtsZ Concentration in *M. gallisepticum* Cells

The concentration estimate (see Figure 6) revealed approximately 150 FtsZ molecules per *M. gallisepticum* cell, corresponding to a concentration of ∼3 µM. This value is consistent with the FtsZ concentration observed in *E. coli* cells (3–10 µM) and exceeds the critical concentration required for *M. gallisepticum* FtsZ polymerization (see below). Thus, the data support the hypothesis that FtsZ can polymerize in *M. gallisepticum* cells

**Figure 6.**
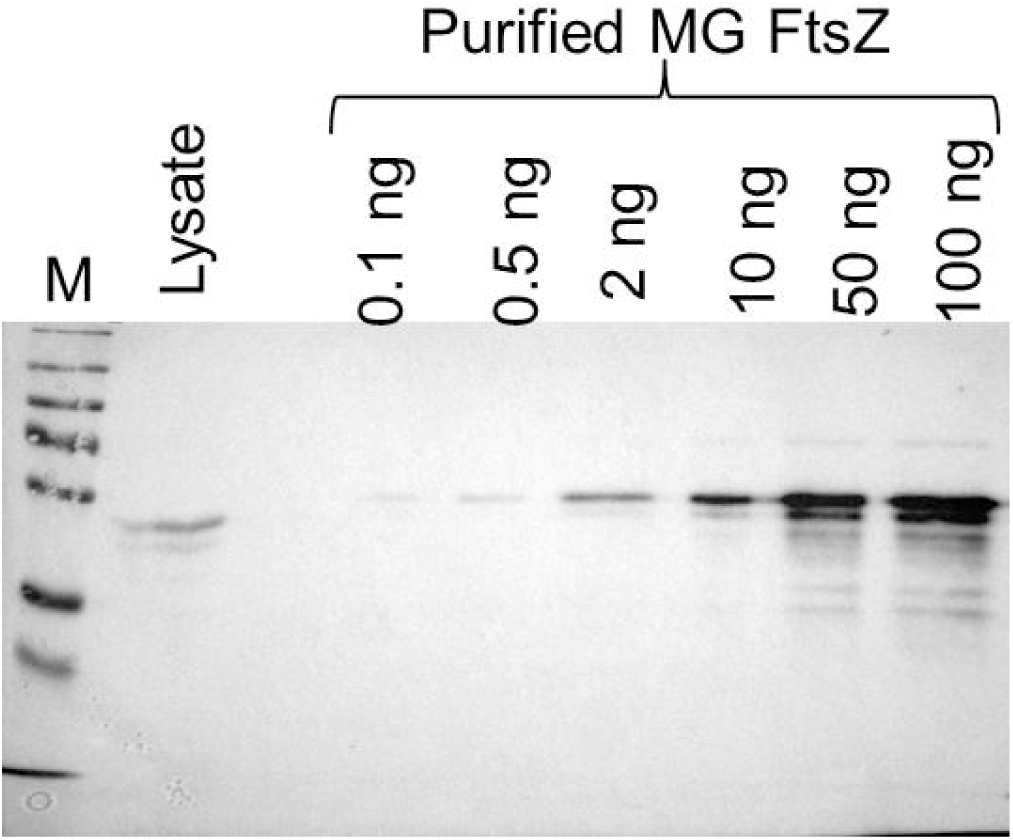
Estimation of FtsZ protein concentration in *M. gallisepticum* cells by semiquantitative immunoblotting. M – molecular weight marker (PageRuler Plus). From left to right: cell lysate from a known number of cells and increasing amounts of purified FtsZ protein.

### 3. Polymerization ability of *M. gallisepticum* FtsZ

The polymerization of *M. gallisepticum* FtsZ in the presence of GTP was analyzed using static light scattering (Figure 7). The protein polymerizes at concentrations above the critical threshold (∼1.2 µM, Figure 7a). Unlike other FtsZ homologs, *M. gallisepticum* FtsZ exhibits slow depolymerization, consistent with its low GTPase activity (discussed later).

**Figure 7.**
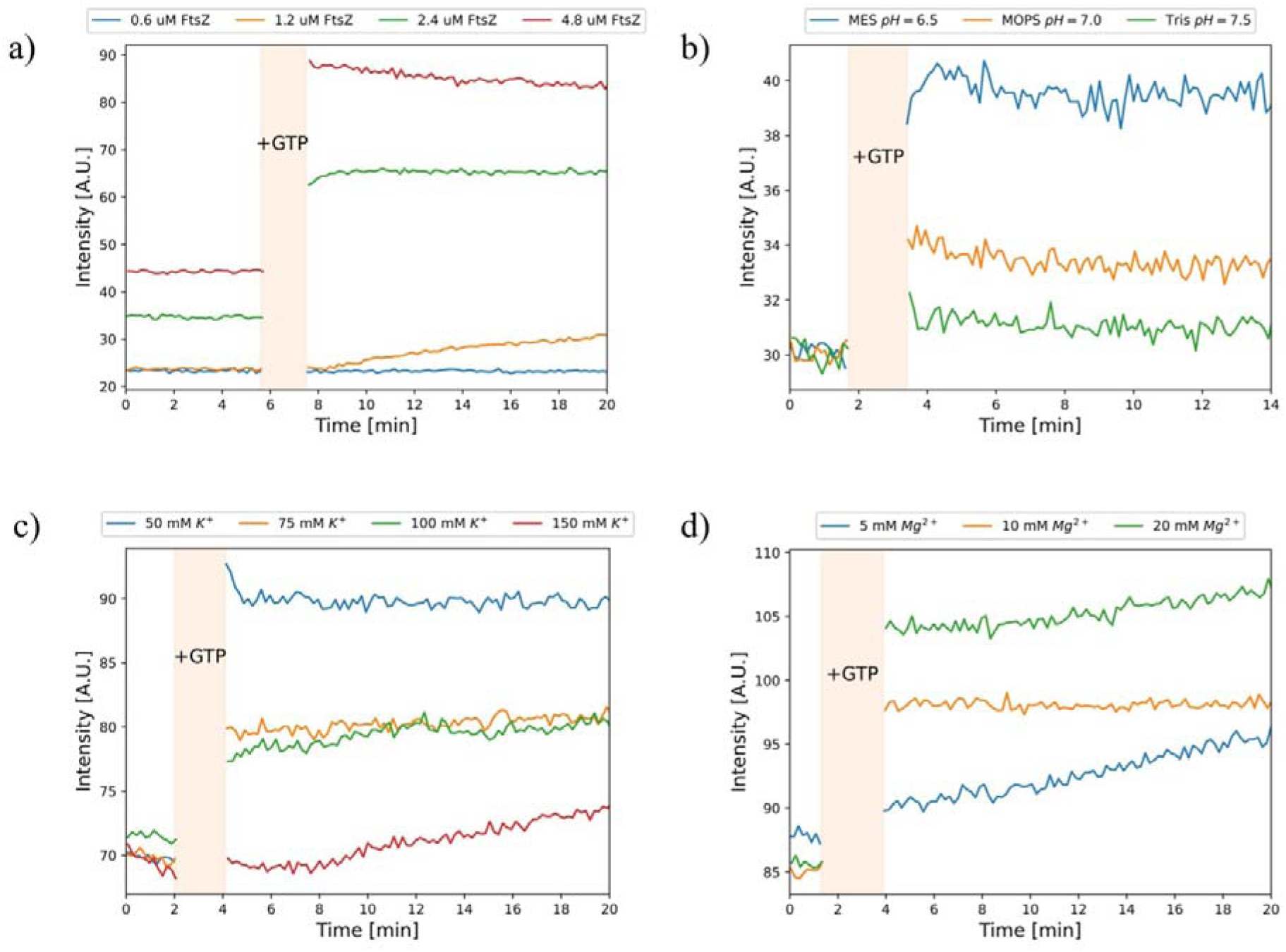
Polymerization curves of the *M. gallisepticum* FtsZ protein. a) Dependence of the polymerization pattern on the FtsZ concentration. MES-based buffers (pH 6.5) were used. The potassium ion concentration in each buffer was 50 mM. The buffers also contained 1 mM EDTA and 5 mM MgCl□. b) Dependence of the polymerization pattern on the pH level for the *M. gallisepticum* FtsZ protein. MES-based (pH 6.5), MOPS (pH 7.0), and Tris (pH 7.5) buffers were used. The potassium ion concentration in each buffer was 50 mM. The buffers also contained 1 mM EDTA and 5 mM MgCl□. c) Dependence of the polymerization pattern on the K□ ion concentration for the *M. gallisepticum* FtsZ protein. MES-based buffers (pH 6.5) were used. The potassium ion concentration was varied by adding KCl. The buffers also contained 1 mM EDTA and 5 mM MgCl□. d) Dependence of the polymerization pattern on the Mg²□ ion concentration for the *M. gallisepticum* FtsZ protein. MES-based buffers (pH 6.5) were used. The K□ ion concentration in each buffer was 50 mM. The MgCl□ concentration was varied as indicated in the figure. The buffers also contained 1 mM EDTA. The beige vertical stripe in the graphs marks the gap corresponding to the addition of GTP.

Effects of environmental factors on polymerization:

- pH: Increased pH inhibits polymerization (Figure 7b), likely due to disruption of electrostatic interactions between monomers in protofilaments [35] and lateral interactions between protofilaments [36].
- Mg² concentration: The optimal MgCl concentration for polymerization is ∼20 mM (Figure 7d). Lower Mg² concentrations significantly reduce polymerization efficiency.
- K concentration: Elevated K levels strongly inhibit polymerization (Figure 7c). At >100 mM K, polymerization is nearly abolished.

In *E. coli* and *B. subtilis*, 100 mM K promotes optimal filament dynamics [37]. *B. subtilis* FtsZ also shows pH-dependent polymerization, with minimal assembly at pH 7.5 and 300 mM K [38]. For *M. gallisepticum* FtsZ, high K likely disrupts electrostatic interactions in dimers, consistent with the general trend that higher ionic strength impairs FtsZ polymerization [35].

Despite intracellular K levels of ∼180 mM in *M. gallisepticum* [39], FtsZ likely polymerizes in vivo because:

1. Its concentration (∼3 µM) exceeds the critical threshold;
2. Microscopy data strongly support polymerization under cellular conditions.

It is important to note that the light scattering technique has a significant drawback: it can detect the formation of large particles (protein multimers) but is incapable of distinguishing between single-stranded polymers, bundles, and aggregates. Thus, direct visualization of the polymers is necessary (see below).

Atomic force microscopy (AFM) data also confirm the ability of *M. gallisepticum* FtsZ to polymerize in the presence of GTP (see Figure 8). Linear, straight FtsZ polymers are clearly visible in the presence of GTP: they align on the mica surface and often stick to each other, forming bundle-like structures (see Figure 8, left). Some aggregate-like structures are also visible, even in the absence of GTP (see Figure 8, right). However, in the presence of GTP, these putative aggregates are more abundant.

**Figure 8.**
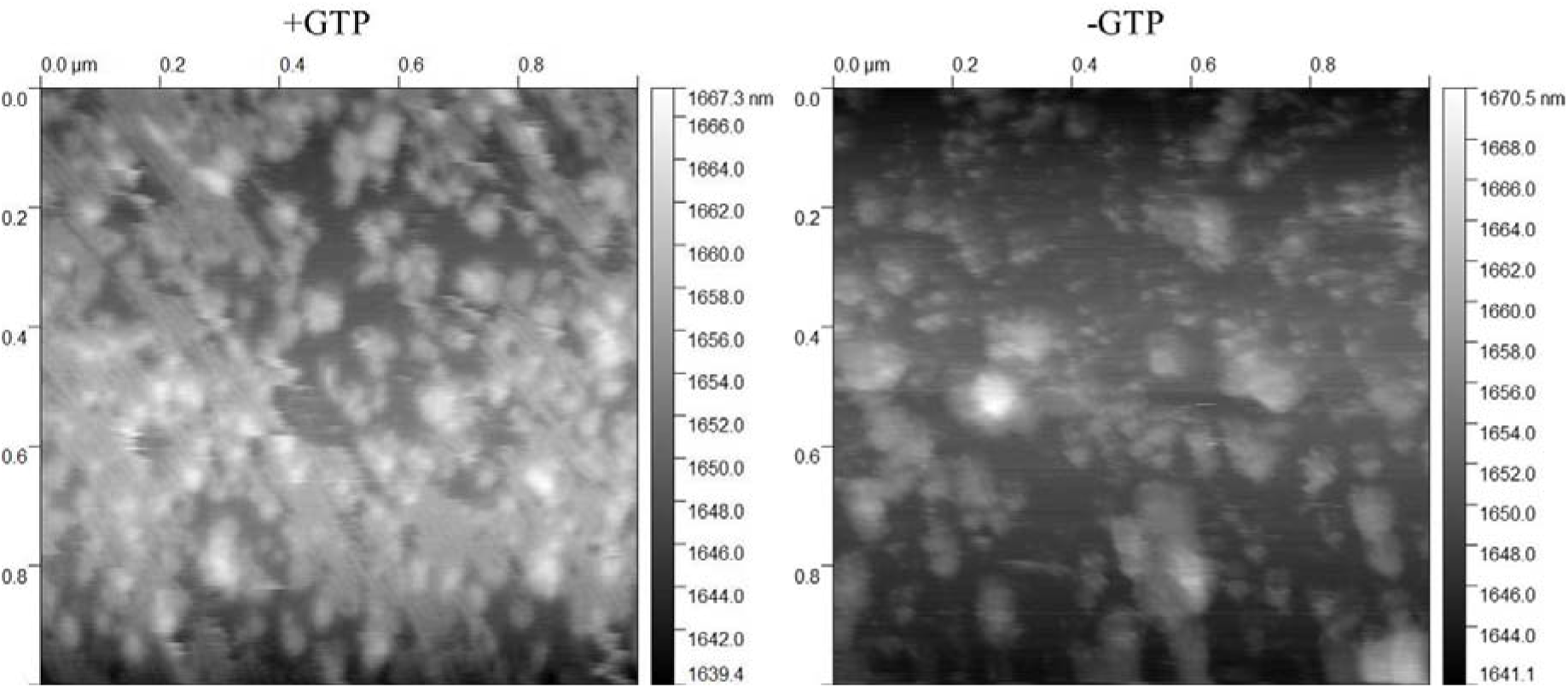
*M. gallisepticum* FtsZ polymers visualized by atomic force microscopy in the presence of GTP (+GTP) and in the absence of GTP (−GTP).

Similar results were obtained using electron microscopy (see Figure S21): *M. gallisepticum* FtsZ forms either bundles or aggregate-like structures. No structures resembling single-stranded FtsZ filaments are visible in the EM images, likely due to the insufficient resolution of the method and/or limitations of the sample preparation protocol. Strikingly, polymerization of *M. gallisepticum* FtsZ is activated not only by GTP but also by ATP (see Figures S21 and S22). This phenomenon is not unique to this protein: it is known that a mutant of *E. coli* FtsZ (FtsZ84) exhibits ATPase activity instead of GTPase activity [40]. We speculate that *M. gallisepticum* FtsZ may function as both a GTPase and an ATPase, but this hypothesis requires further validation.

### 4. GTPase activity of *M. gallisepticum* FtsZ

In this study, the GTPase activity of *M. gallisepticum* FtsZ was measured (Figure 9). On average, one FtsZ molecule hydrolyzes 0.75 ± 0.12 molecules of GTP per minute. This value is approximately 2–3 times lower than that of *E. coli* FtsZ [41] which is consistent with the observed low depolymerization rate of *M. gallisepticum* FtsZ.

**Figure 9.**
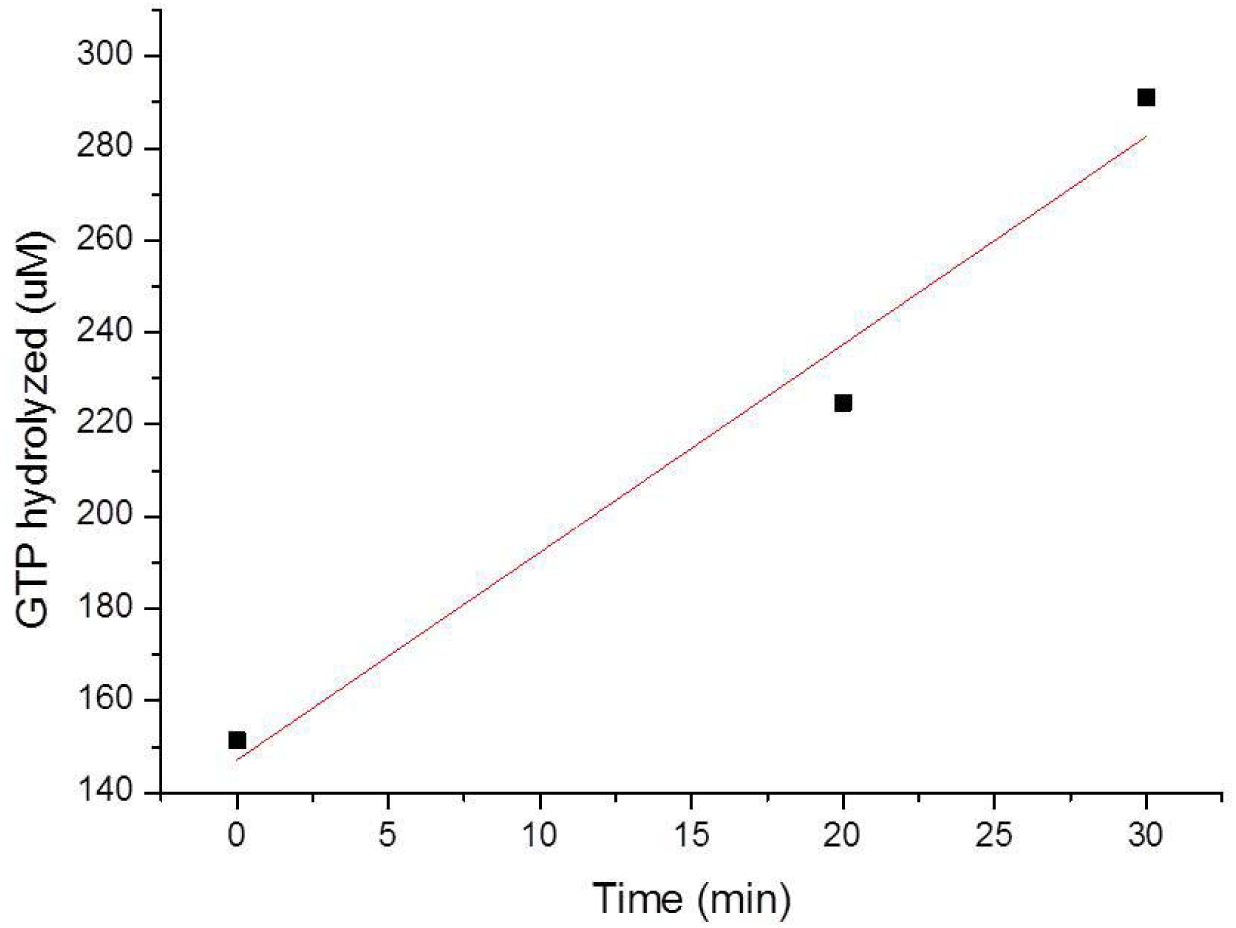
GTPase activity of *M. gallisepticum* FtsZ as measured by the malachite green assay.

### 3. FtsZ Interactions with Other Proteins of *M. gallisepticum*

To more reliably interpret the structures formed by FtsZ in *M. gallisepticum* cells and to better understand its role in mycoplasma physiology, potential FtsZ-interacting proteins were identified in this study. Protein-protein interaction analysis (using co-immunoprecipitation with antibodies against *M. gallisepticum* FtsZ; see Table 1) revealed that FtsZ interacts with several proteins, including elongation factor EF-Tu, lipoprotein A, ATP synthase, the OppA transporter protein, and RNA polymerase.

**Table 1.**
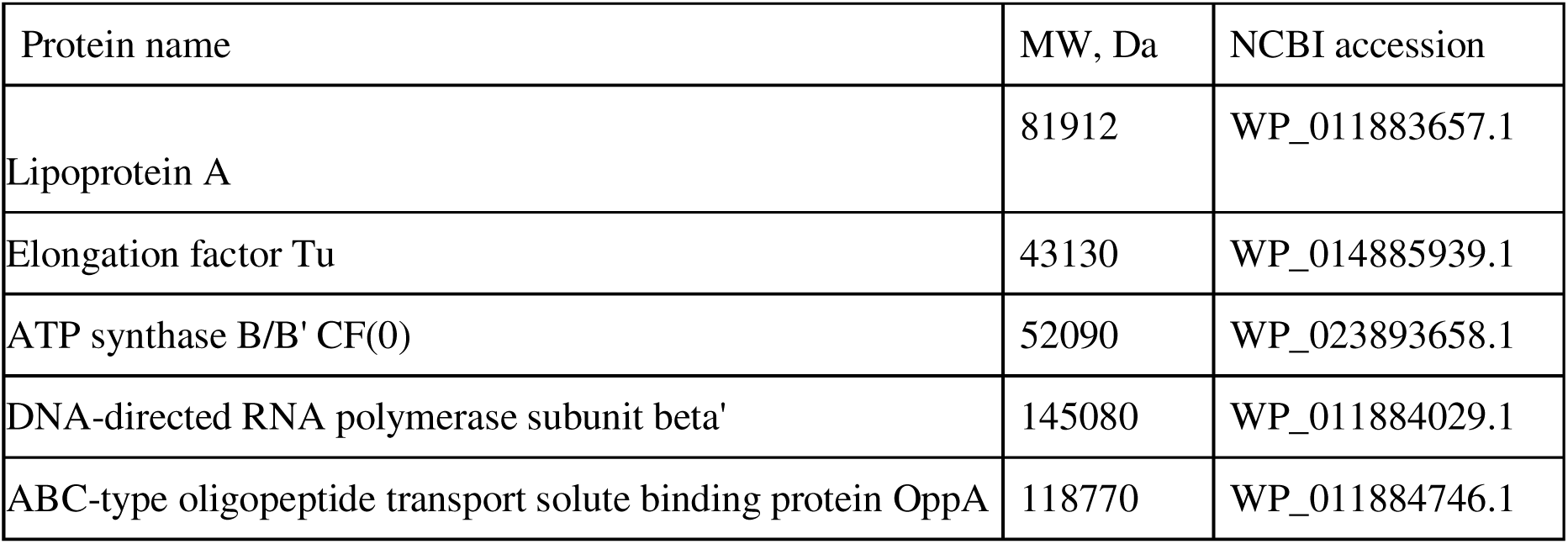
*M. gallisepticum* proteins interacting with FtsZ, as determined by co-immunoprecipitation. MW: molecular weight.

Notably, among the identified interactors, no known homologs of canonical bacterial division proteins (such as FtsK or FtsA) were detected, despite their presence in *M. gallisepticum* and their established interactions with FtsZ in other bacteria. While this observation does not confirm FtsZ’s involvement in cell division, it does not rule it out either, as the absence of these interactions could stem from methodological limitations.

However, some of the detected interactions may provide insights into FtsZ’s potential role in the mycoplasma cytoskeleton. For instance, EF-Tu—a translation elongation factor — has been reported to interact with other cytoskeletal proteins, including MreB [42] and FtsZ [43], and can form filamentous structures *in vitro* [44]. The observed interaction between FtsZ and EF-Tu suggests their possible cooperation in forming cytoskeletal structures in mycoplasma cells.

To further characterize protein–protein interactions involving *M. gallisepticum* FtsZ, a bacterial two-hybrid system was used [45]. Several proteins were selected as putative partners of FtsZ: EF-Tu (based on co-immunoprecipitation data), FtsK and FtsA (based on homology with other organisms), and GapD (identified as an FtsZ partner in some co-immunoprecipitation replicates in our study). Additionally, the FtsZ-FtsZ interaction was analyzed as a positive control. The results are shown in Figure 10 and Table 2.

**Figure 10.**
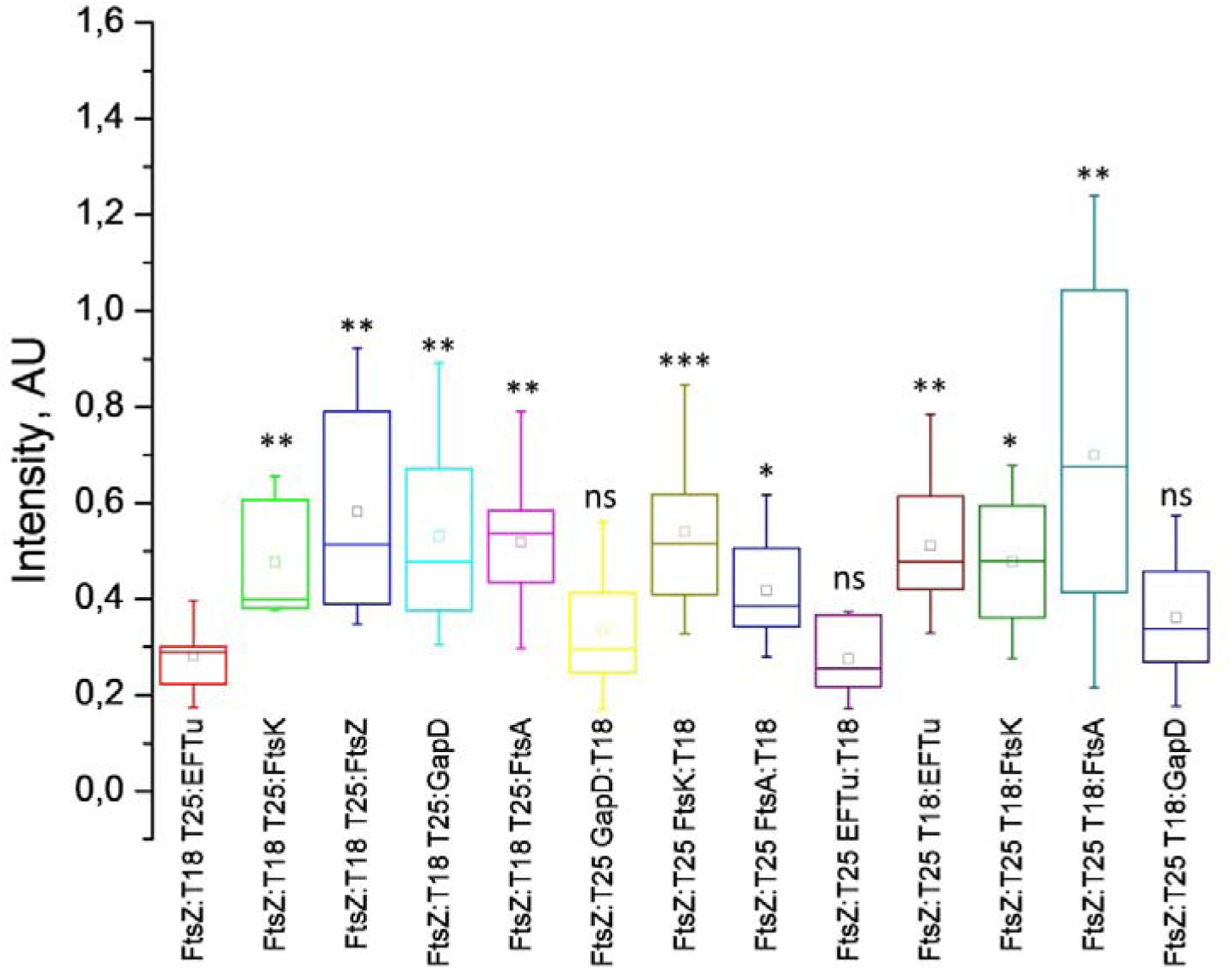
Comparison of the signal intensity of β-galactosidase activity, reflecting protein-protein interactions between the specified protein pairs. The box-and-whisker plots show the quantile distributions of activity values across replicates: the box boundaries represent the 2nd and 3rd quartiles, while the whiskers indicate the 5th and 95th percentiles. The square inside the box denotes the mean value, and the horizontal line represents the median. Asterisks indicate the significance levels for differences in value distributions compared to the FtsZ:T18 + T25:EF-Tu sample (one of the samples with the lowest signal): * – significance level p<0.05; ** – significance level p<0.01; *** – significance level p<0.001. Pairwise comparisons were performed using the Mann-Whitney test.

**Table 2.** Median and p-value for for β-galactosidase activity signal intensities, reflecting protein-protein interactions between the specified protein pairs.

| Protein pair: | FtsZ:T18 T25:EFTu | FtsZ:T18 T25:FtsK | FtsZ:T18 T25:FtsZ | FtsZ:T18 T25:GapD | FtsZ:T18 T25:FtsA | FtsZ:T25 GapD:T18 | FtsZ:T25 FtsK:T18 | FtsZ:T25 FtsA:T18 | FtsZ:T25 EFTu:T18 | FtsZ:T25 T18:EFTu | FtsZ:T25 T18:FtsK | FtsZ:T25 T18:FtsA | FtsZ:T25 T18:GapD |
| --- | --- | --- | --- | --- | --- | --- | --- | --- | --- | --- | --- | --- | --- |
| Median | 0,29 | 0,40 | 0,51 | 0,48 | 0,54 | 0,30 | 0,51 | 0,39 | 0,26 | 0,48 | 0,48 | 0,68 | 0,34 |
| p-value | - | 0,0030 | 0,0031 | 0,0021 | 0,0021 | 0,6065 | 0,0003 | 0,0164 | 0,6065 | 0,0012 | 0,0140 | 0,0079 | 0,4079 |

The interactions of *M. gallisepticum* FtsZ with EF-Tu, FtsA, FtsK, and GapD were confirmed. The observed interactions with FtsA and FtsK support the involvement of FtsZ in the cell division process of *M. gallisepticum*.

### Conclusions

1. Unlike well-studied bacteria, the FtsZ in *M. gallisepticum* shows polar rather than mid-cell localization.
2. Overexpression of fluorescently labeled FtsZ enhances its polar localization and may lead to mini-cell formation.
3. The FtsZ concentration in the cell was measured (approximately 3 μM) and, together with *in vitro* data (confirming its ability to form polymers and hydrolyze GTP), supports its ability to polymerize *in vivo*.
4. *In vitro* studies confirm that *M. gallisepticum* FtsZ can polymerize and hydrolyze GTP. Its GTPase activity is relatively low, and polymers are disassembled slowly.
5. Protein-protein interactions between *M. gallisepticum* FtsZ and FtsA, FtsK, EF-Tu, and GapD were identified. Interaction with FtsA and FtsK supports the involvement of FtsZ in *M. gallisepticum* cell division.

Overall, the results support the role of FtsZ in *M. gallisepticum* cell division, though its properties significantly differ from those of other known homologs.

It would be interesting to determine whether FtsZ is essential for *M. gallisepticum* cell division, as the obtained evidence suggests its participation in the process. Interestingly, a Tn4001 transposon random insertion into the *ftsZ* gene was not lethal and did not cause growth inhibition in *M. gallisepticum* cells (unpublished data; the strain was isolated by G. Yu. Fisunov and kindly provided to us). Thus, we hypothesize that FtsZ is not essential for *M. gallisepticum* cell division, possibly due to the presence of multiple mechanisms with overlapping functions (e.g., FtsZ-dependent and motility-dependent). However, further studies are needed to confirm this hypothesis.

Notably, in another motile mycoplasma, *M. genitalium*, FtsZ is also nonessential for viability, as the cells can divide via binary fission, presumably mediated by gliding motility, which depends on proteins of the terminal organelle and associated cytoskeletal structures [13]. Interestingly, mollicutes, which may employ different division modes, have retained the *ftsZ* gene. For example, in *M. genitalium*, FtsZ appears to be the primary driver of division during host cell infection [13]. In contrast, the division mechanism of mollicutes lacking the *ftsZ* gene remains unknown.

## Declaration of generative AI in scientific writing

During the preparation of this work, the authors used DeepSeek-V3 to correct grammatical errors, fix typos, and improve the text’s readability. After applying this tool, the authors reviewed and edited the content as necessary and take full responsibility for the publication’s content.

## Acknowledgments

The study was supported by grant of the Russian Science Foundation No. 24-74-10022, https://rscf.ru/project/24-74-10022/. The work was carried out using scientific equipment of the Center of Shared Usage “The Analytical Center of Nano- and Biotechnologies of SPbPU”. The authors are grateful to G.Y. Fisunov for providing the *M. gallisepticum* S6 wild-type strain, the *ftsZ* knockout strain, and the pTn4001mini_Recipient-MCS-M5 vector. The authors also thank A.R. Kayumov for supplying the vectors for the bacterial two-hybrid system and T.P. Sankova for providing the K802 strain. Special thanks go to N.E. Morozova for carefully reading the manuscript and offering valuable advice.

## Author Contributions

N.R.: Investigation, Formal analysis; D.G.: Investigation, Formal analysis; A.K: Investigation, Formal analysis; A.S.: Investigation, Formal analysis; Y.B.: Investigation, Formal analysis; E.P.: Investigation, Formal analysis; V.P.: Investigation, Formal analysis; A.D.: Formal analysis; V.I: Investigation, Formal analysis; T.A: Investigation, Formal analysis; A.A: Investigation, Formal analysis; I.V.: Investigation, Formal analysis; M.K: Formal analysis; A.V.: Investigation, Formal analysis, Supervision, Writing, Funding acquisition.

