## Supplementary materials for "*Mycoplasma gallisepticum* FtsZ demonstrates properties that distinguish it from other known homologs"

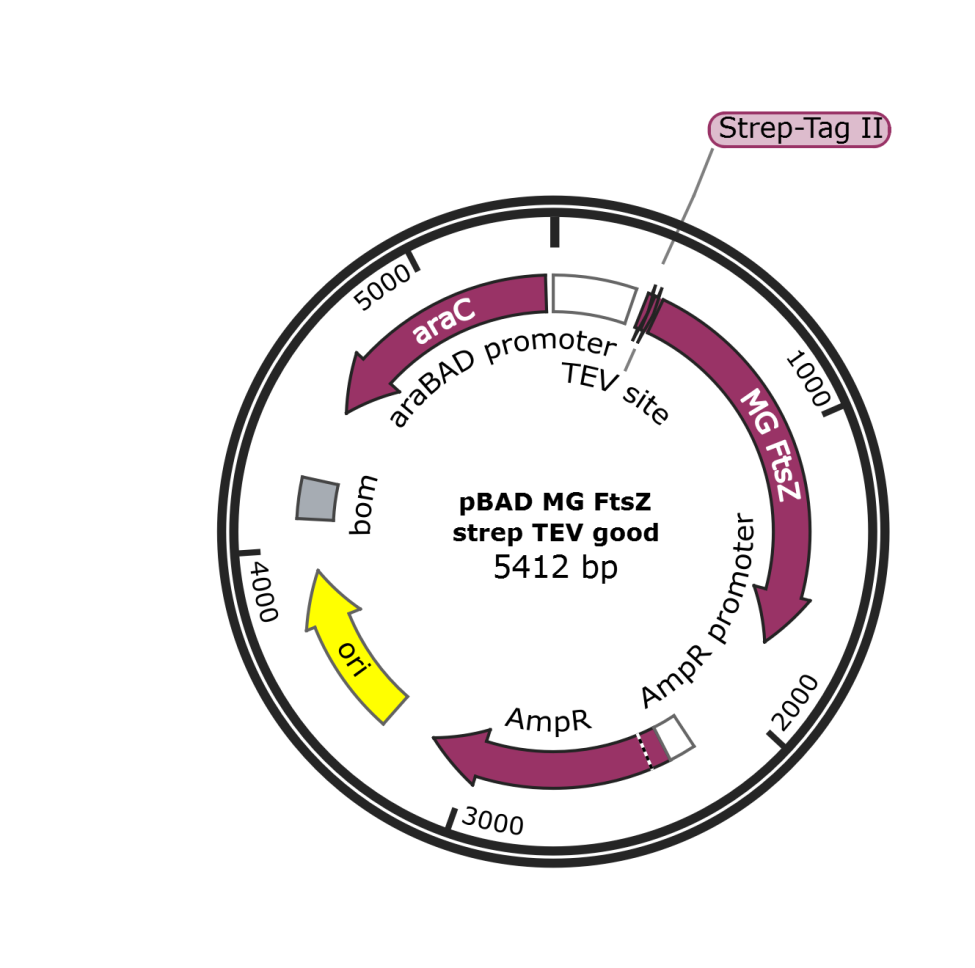

**Figure S1 –** Map of the plasmid used for *M. gallisepticum* FtsZ purification for immunization. The main elements of the plasmid are shown in the figure. Sequence of the plasmid is shown in file “pBAD MG FtsZ strep TEV good.gb”

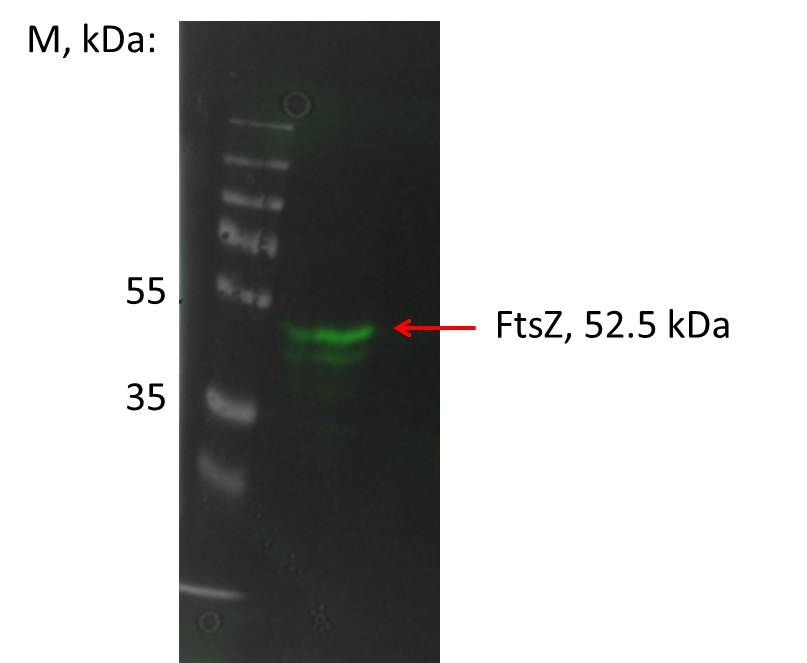

**Figure S2 –** Evaluation of the specificity of antibodies to the FtsZ protein by Western-blotting. A combined image based on the membrane image in transmitted light (grayscale) and the image in the chemiluminescence channel (green) is shown. One major band is visible on the membrane, approximately the size of the target protein (52.5 kDa)

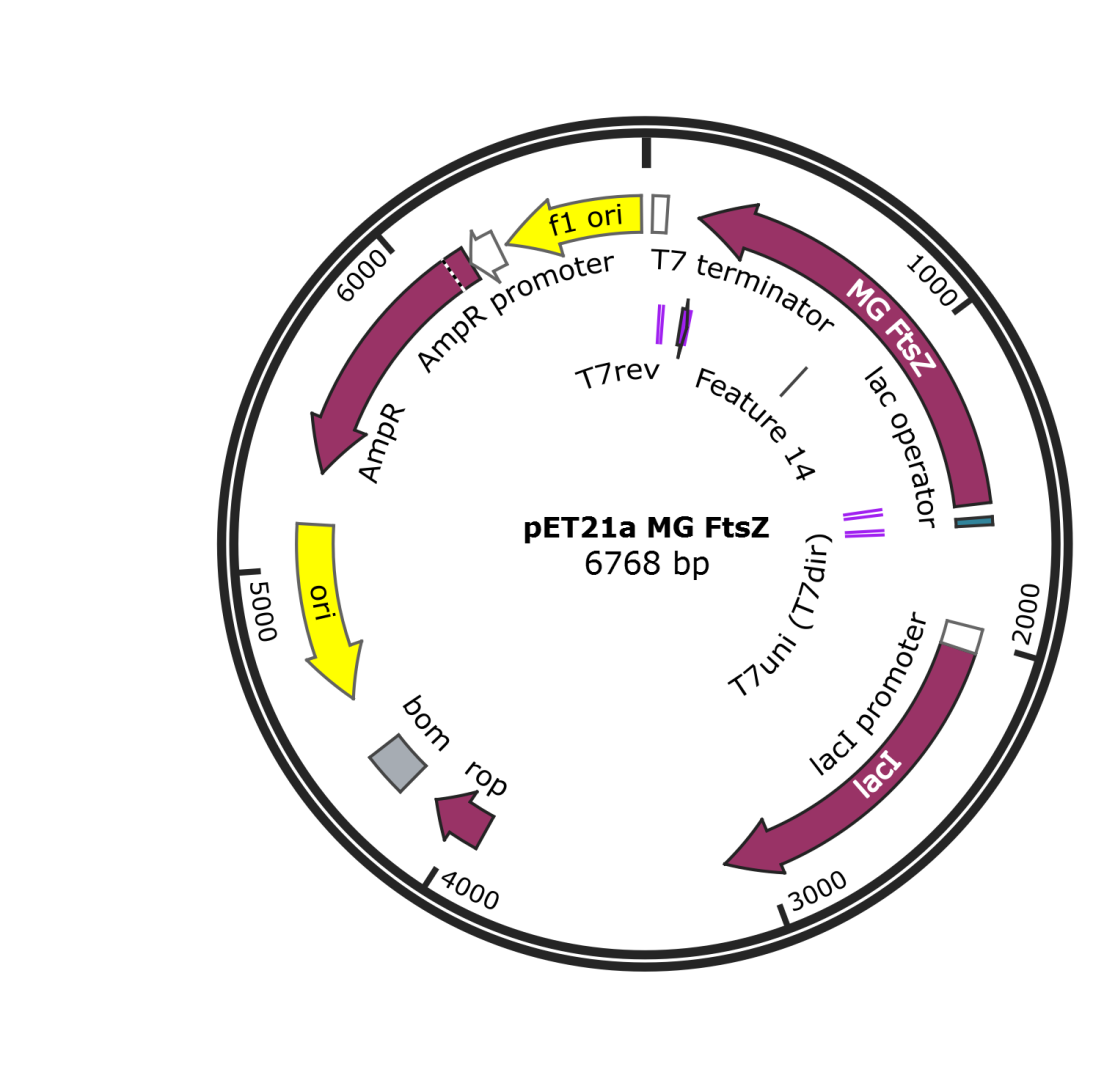

**Figure S3 –** Map of the plasmid used for *M. gallisepticum* FtsZ purification for *in vitro* experiments (polymerization and GTPase activity). The main elements of the plasmid are shown in the figure. Sequence of the plasmid is shown in file “pET21a MG FtsZ.gb”

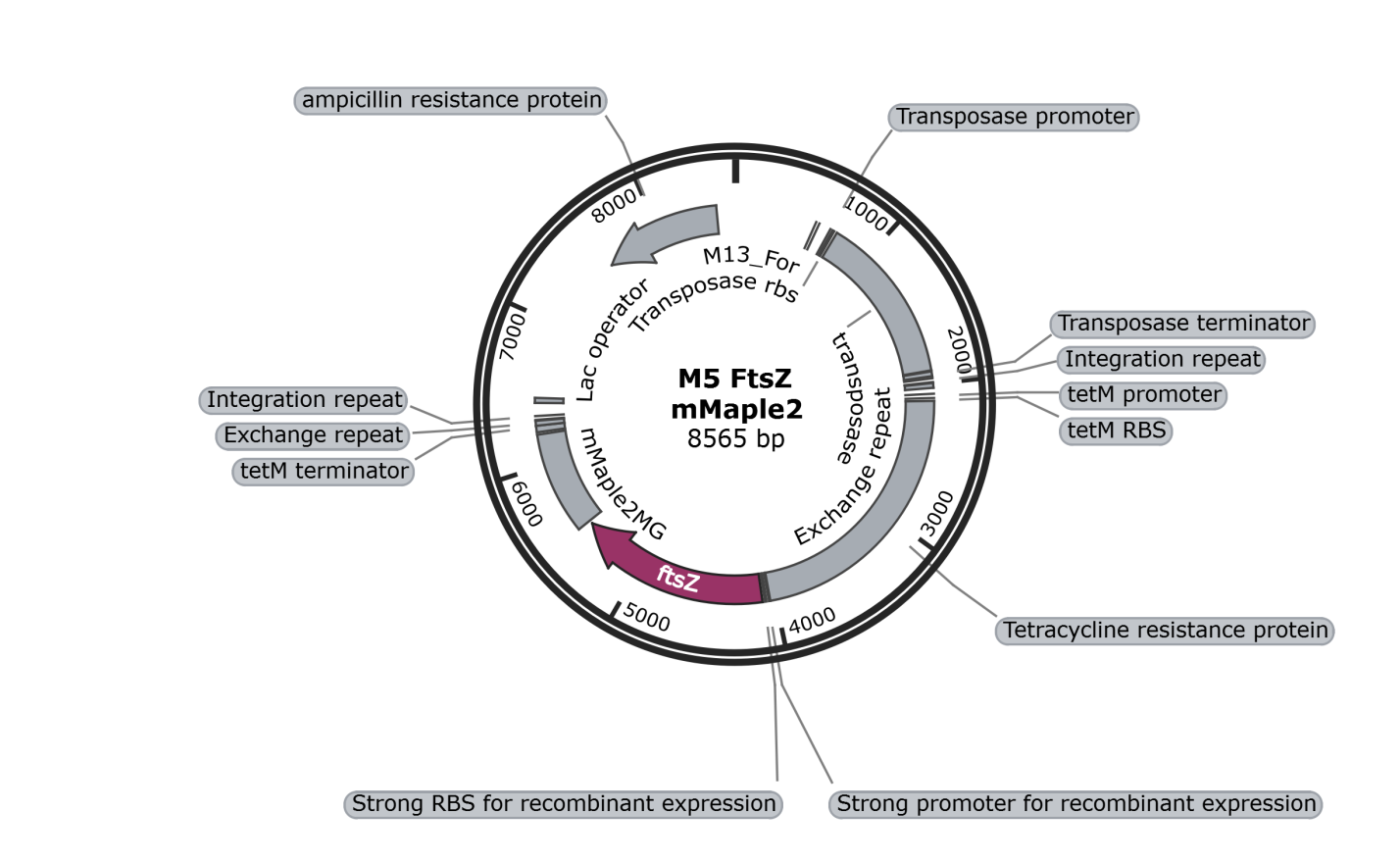

**Figure S4 –** Map of the plasmid used for TN4001 transposon mutagenesis of *M. gallisepticum* to obtain a strain overproducing FtsZ:mMaple2 fusion. The main elements of the plasmid are shown in the figure. Sequence of the plasmid is shown in file “M5 FtsZ mMaple2.gb”

**Table S1** – Maps of plasmids used for bacterial two-hybrid system to study interactions of FtsZ with other proteins of *M. gallisepticum*

| **Plasmid** | **Figure name** | **File name** |
| --- | --- | --- |
| FtsZ:T18 | Figure S5 | FtsZ-T18 Map.gb |
| FtsZ:T25 | Figure S6 | FtsZ-T25 Map.gb |
| T25:EFTu | Figure S7 | T25-EFTu Map.gb |
| T25:FtsA | Figure S8 | T25-FtsA Map.gb |
| T25:FtsK | Figure S9 | T25-FtsK Map.gb |
| T25:FtsZ | Figure S10 | T25-FtsZ Map.gb |
| T25:GapD | Figure S11 | T25-GapD Map.gb |
| EFTu:T18 | Figure S12 | EFTu-T18 Map.gb |
| FtsA:T18 | Figure S13 | FtsA-T18 Map.gb |
| FtsK:T18 | Figure S14 | FtsK-T18 Map.gb |
| GapD:T18 | Figure S15 | GapD-T18 Map.gb |
| T18:EFTu | Figure S16 | T18-EFTu Map.gb |
| T18:FtsA | Figure S17 | T18-FtsA Map.gb |
| T18:FtsK | Figure S18 | T18-FtsK Map.gb |
| T18:GapD | Figure S19 | T18-GapD Map.gb |

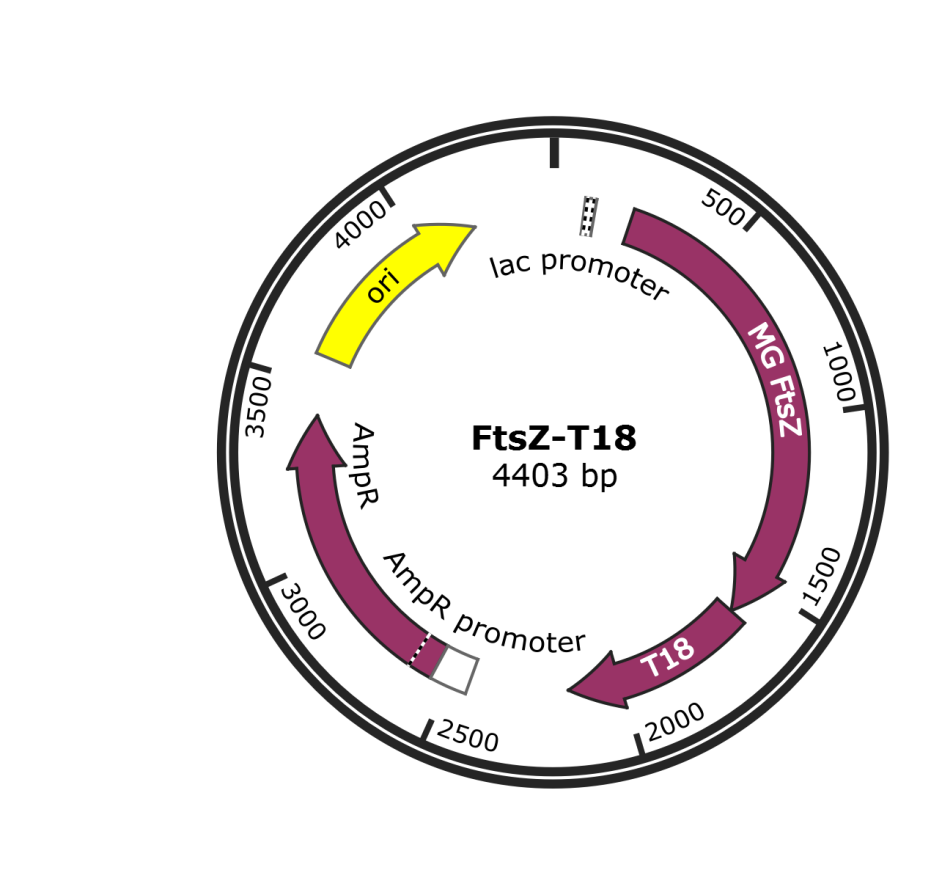

**Figure S5 –** Map of the plasmid FtsZ-T18 used for analysis of *M. gallisepticum* FtsZ interactions with other proteins. The main elements of the plasmid are shown in the figure. Sequence of the plasmid is shown in file “FtsZ-T18.gb”

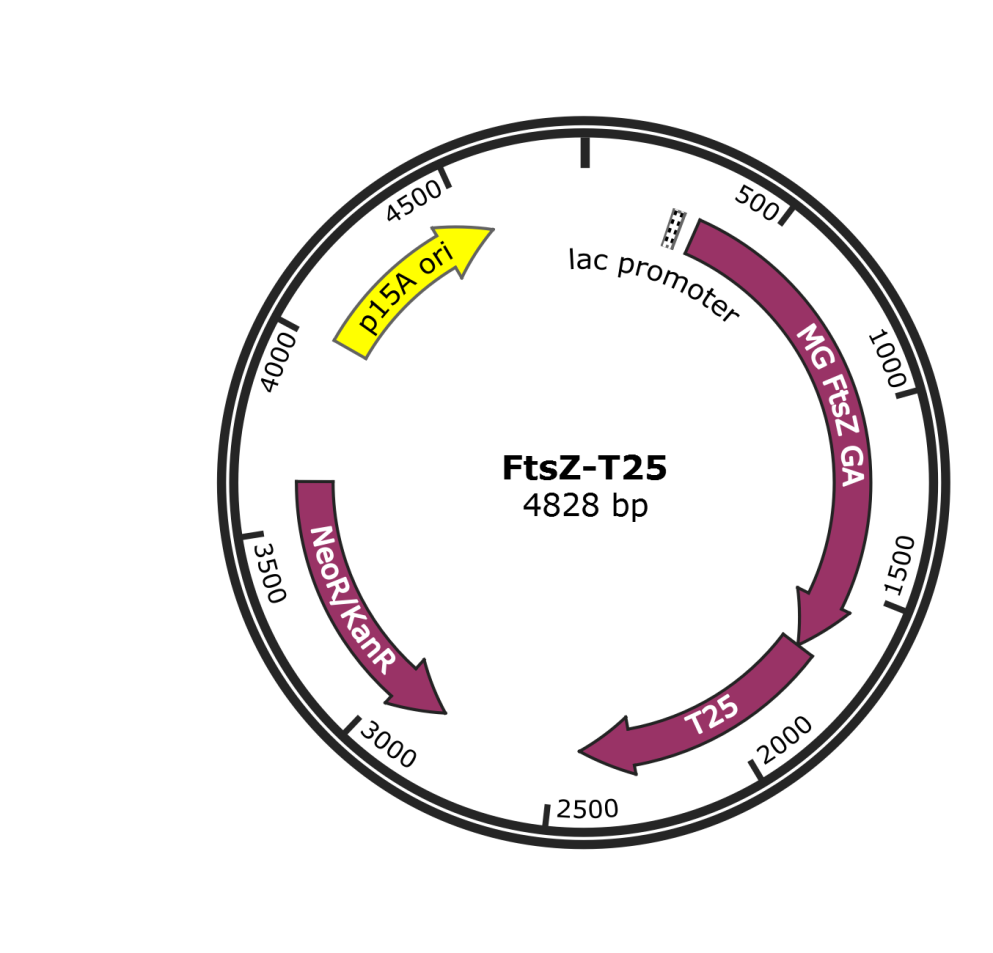

**Figure S6 –** Map of the plasmid FtsZ-T25 used for analysis of *M. gallisepticum* FtsZ interactions with other proteins. The main elements of the plasmid are shown in the figure. Sequence of the plasmid is shown in file “FtsZ-T25.gb”

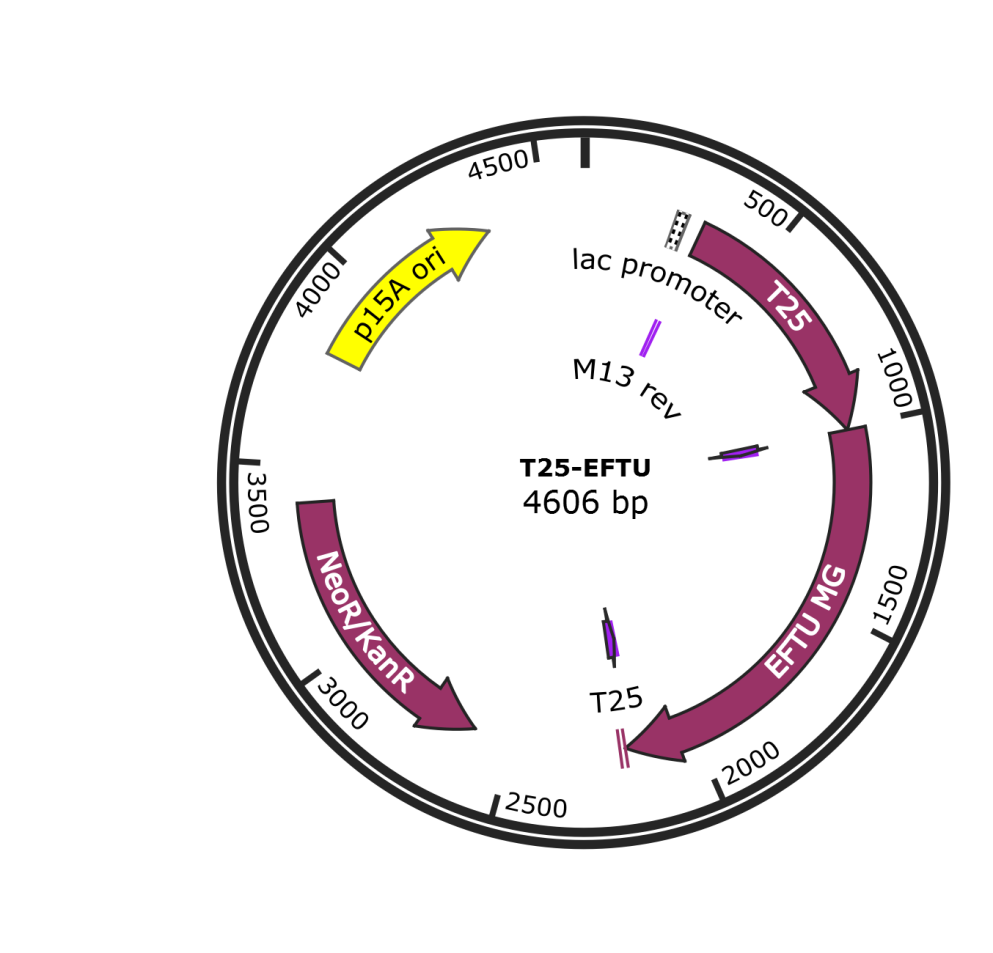

**Figure S7 –** Map of the plasmid T25-EFTU used for analysis of *M. gallisepticum* FtsZ interactions with other proteins. The main elements of the plasmid are shown in the figure. Sequence of the plasmid is shown in file “T25-EFTU.gb”

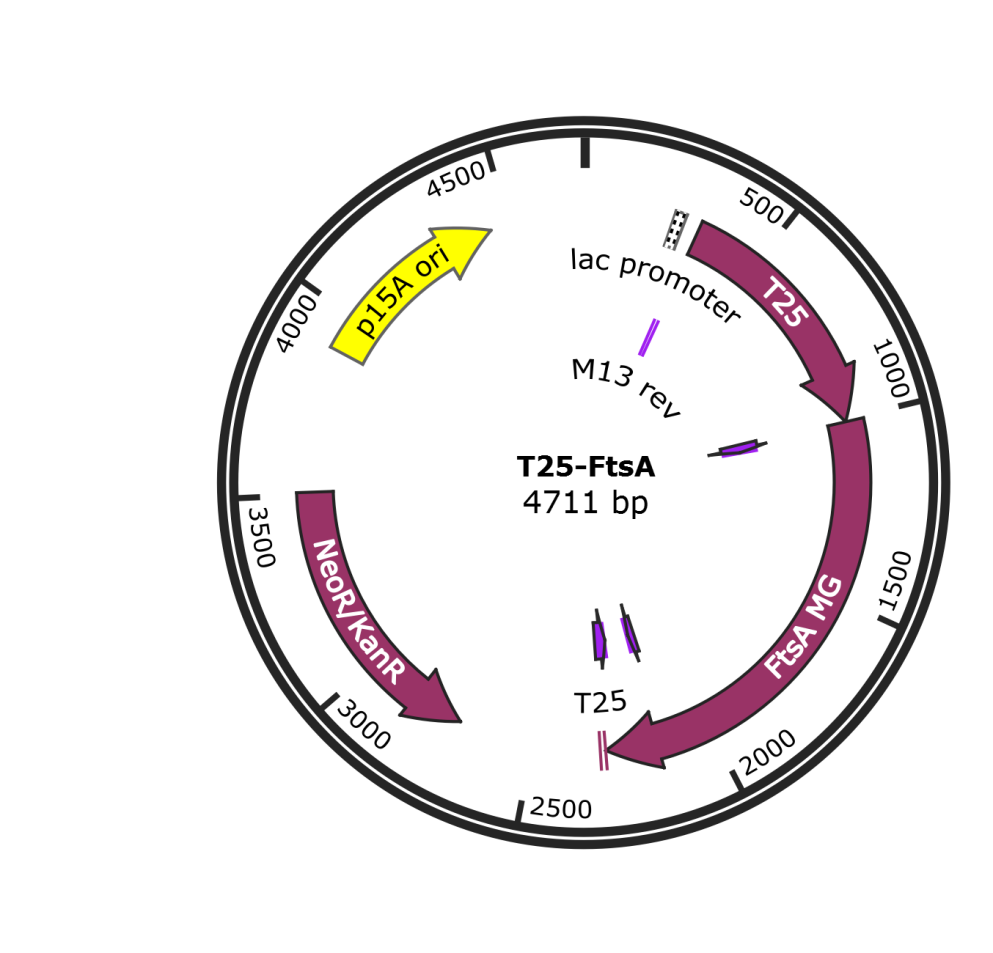

**Figure S8 –** Map of the plasmid T25-FtsA used for analysis of *M. gallisepticum* FtsZ interactions with other proteins. The main elements of the plasmid are shown in the figure. Sequence of the plasmid is shown in file “T25-FtsA.gb”

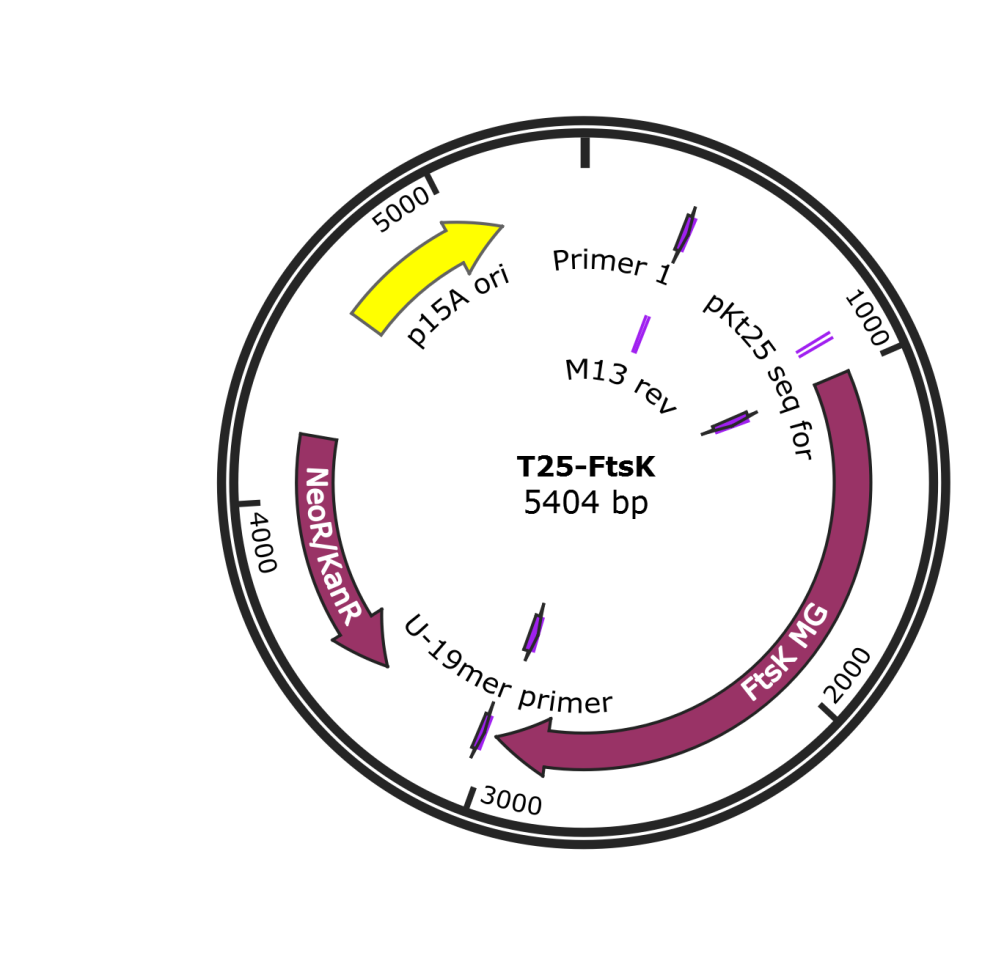

**Figure S9 –** Map of the plasmid T25-FtsK used for analysis of *M. gallisepticum* FtsZ interactions with other proteins. The main elements of the plasmid are shown in the figure. Sequence of the plasmid is shown in file “T25-FtsK.gb”

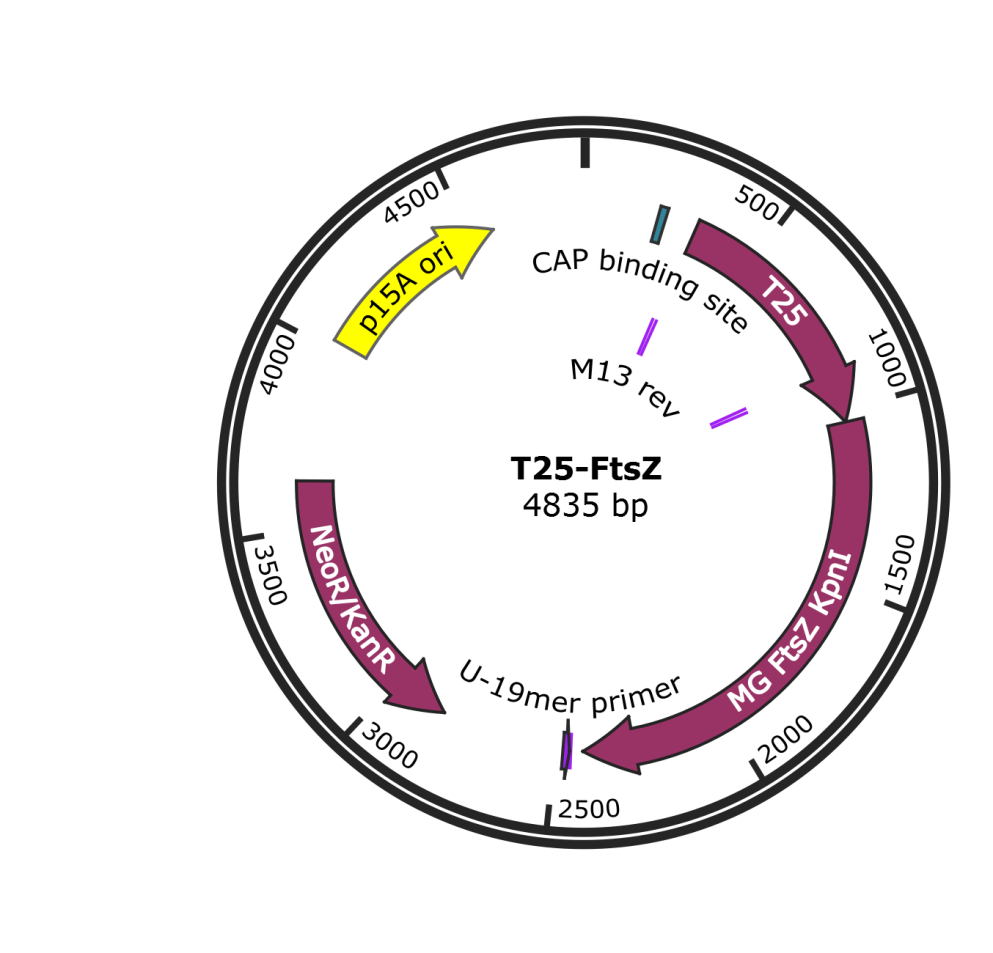

**Figure S10 –** Map of the plasmid T25-FtsZ used for analysis of *M. gallisepticum* FtsZ interactions with other proteins. The main elements of the plasmid are shown in the figure. Sequence of the plasmid is shown in file “T25-FtsZ.gb”

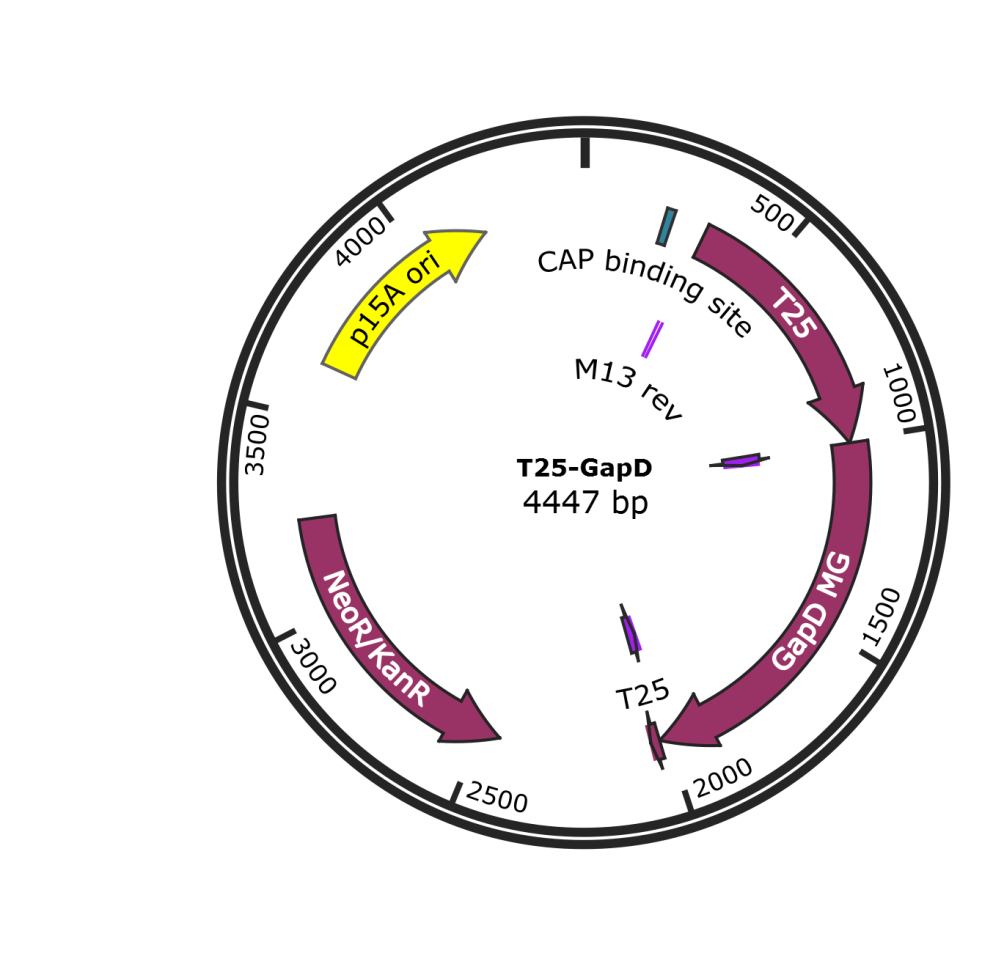

**Figure S11 –** Map of the plasmid T25-GapD used for analysis of *M. gallisepticum* FtsZ interactions with other proteins. The main elements of the plasmid are shown in the figure. Sequence of the plasmid is shown in file “T25-GapD.gb”

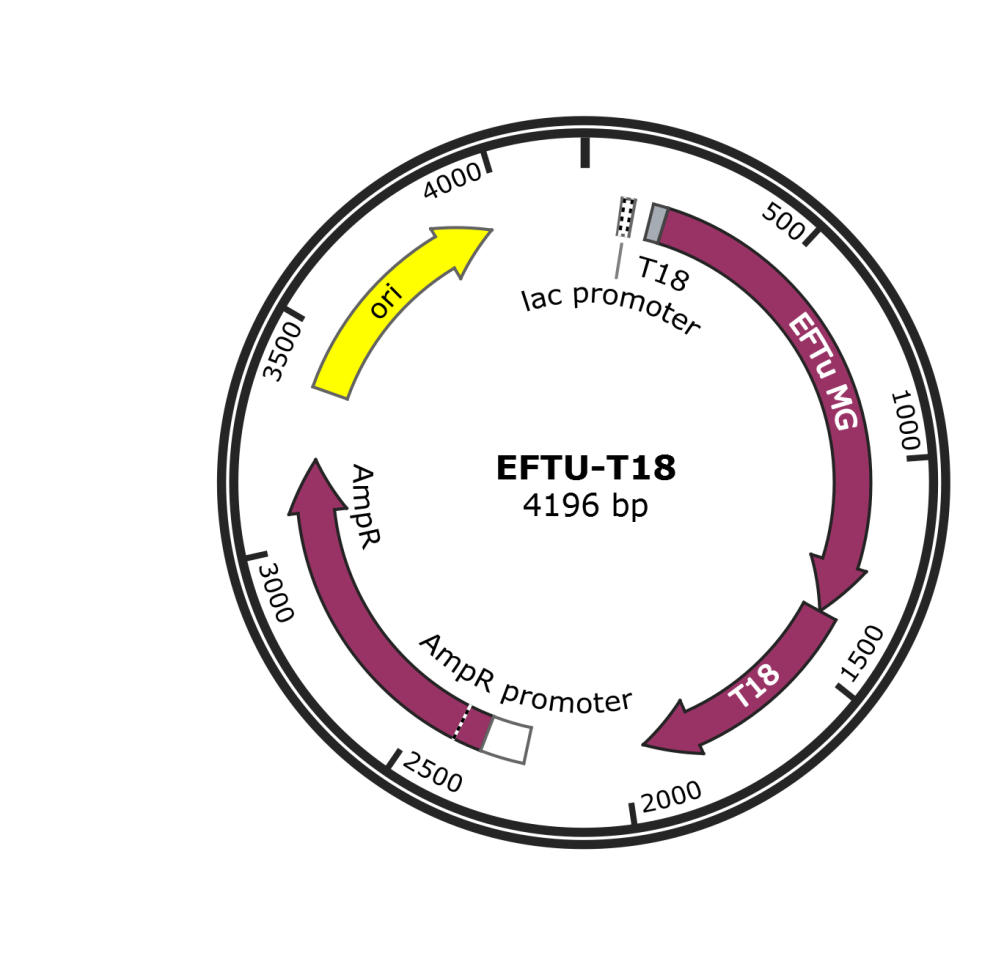

**Figure S12 –** Map of the plasmid EFTU-T18 used for analysis of *M. gallisepticum* FtsZ interactions with other proteins. The main elements of the plasmid are shown in the figure. Sequence of the plasmid is shown in file “EFTU-T18.gb”

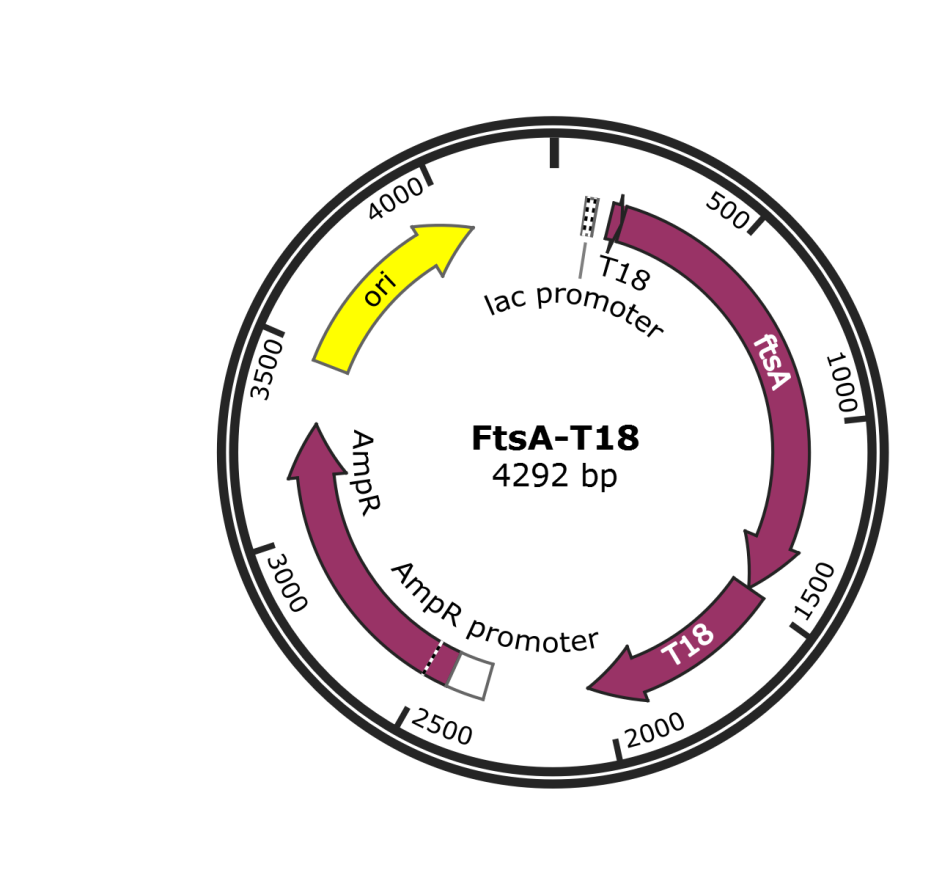

**Figure S13 –** Map of the plasmid FtsA-T18 used for analysis of *M. gallisepticum* FtsZ interactions with other proteins. The main elements of the plasmid are shown in the figure. Sequence of the plasmid is shown in file “FtsA-T18.gb”

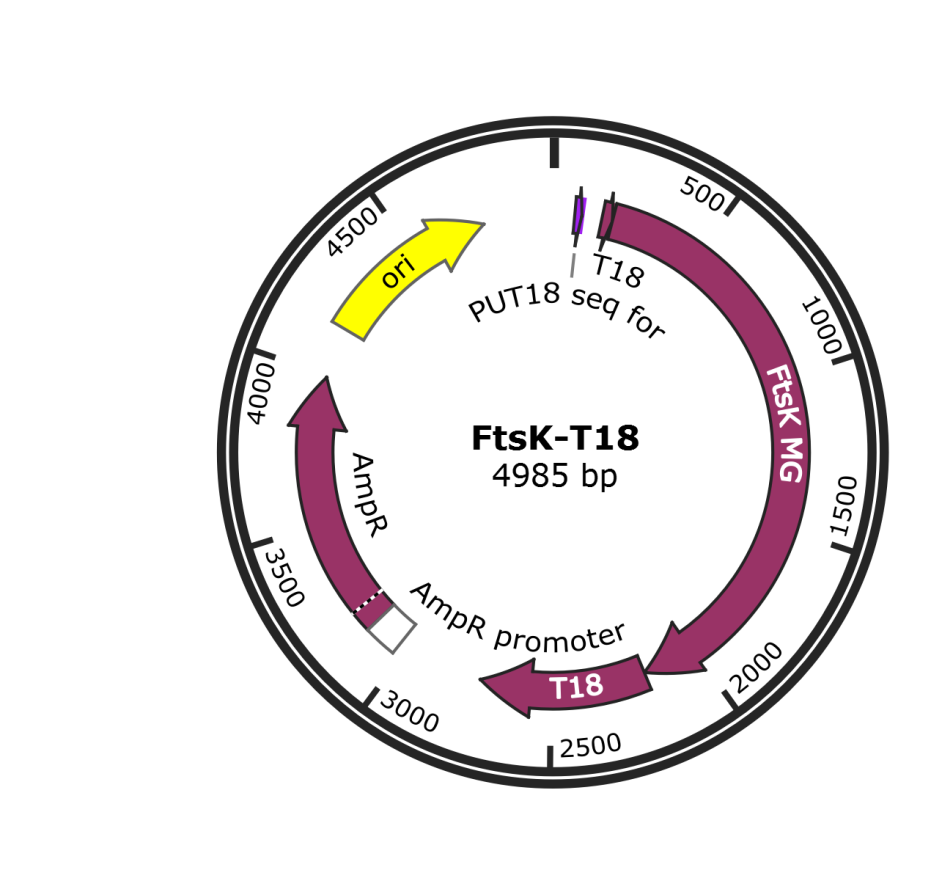

**Figure S14 –** Map of the plasmid FtsK-T18 used for analysis of *M. gallisepticum* FtsZ interactions with other proteins. The main elements of the plasmid are shown in the figure. Sequence of the plasmid is shown in file “FtsK-T18.gb”

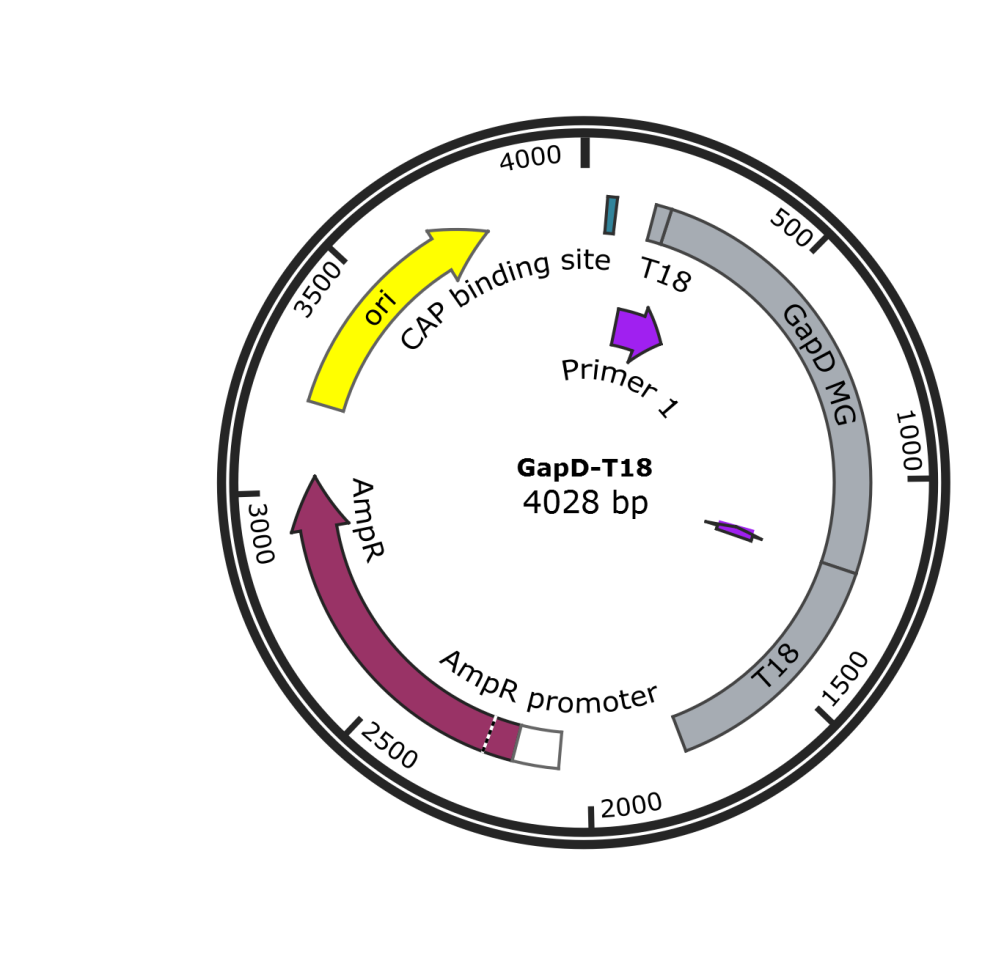

**Figure S15 –** Map of the plasmid GapD-T18 used for analysis of *M. gallisepticum* FtsZ interactions with other proteins. The main elements of the plasmid are shown in the figure. Sequence of the plasmid is shown in file “GapD-T18.gb”

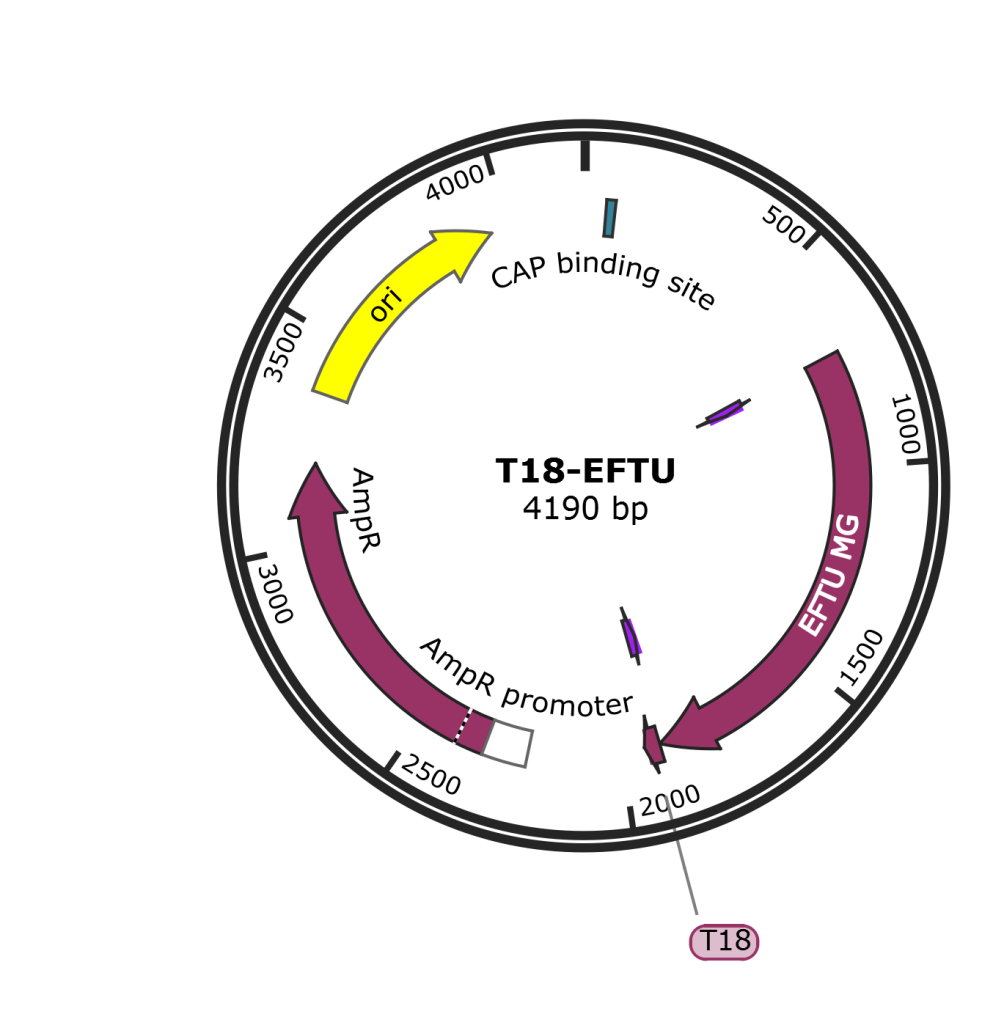

**Figure S16 –** Map of the plasmid T18-EFTU used for analysis of *M. gallisepticum* FtsZ interactions with other proteins. The main elements of the plasmid are shown in the figure. Sequence of the plasmid is shown in file “T18-EFTU.gb”

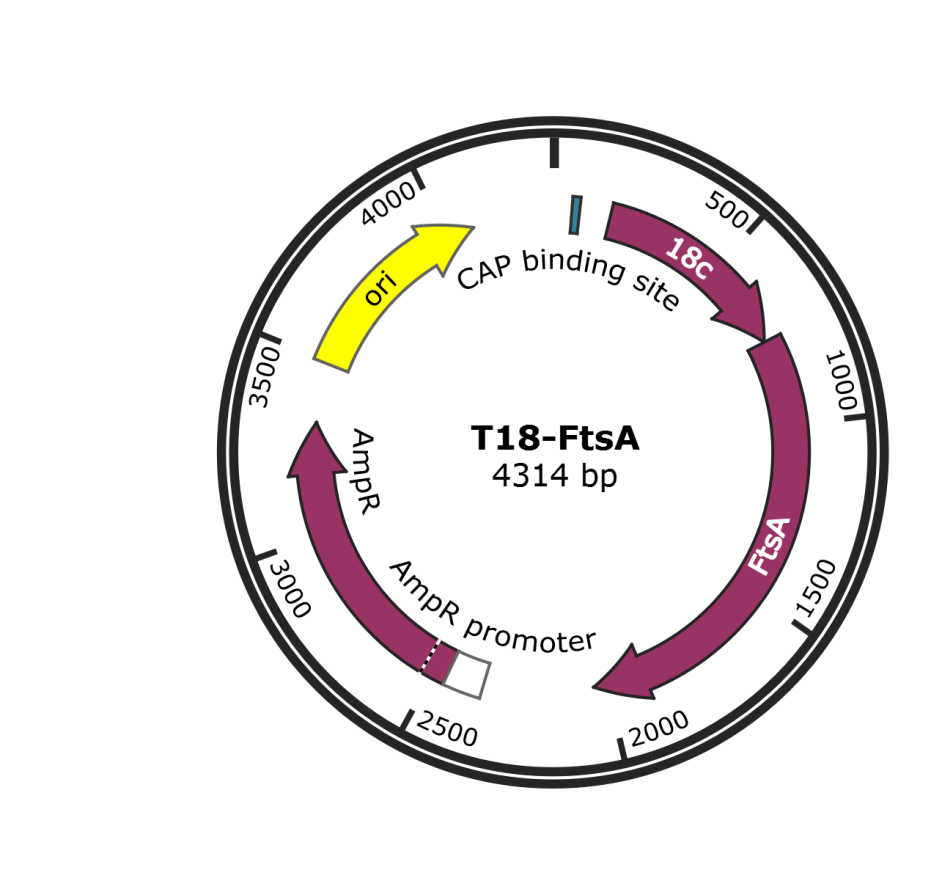

**Figure S17 –** Map of the plasmid T18-FtsA used for analysis of *M. gallisepticum* FtsZ interactions with other proteins. The main elements of the plasmid are shown in the figure. Sequence of the plasmid is shown in file “T18-FtsA.gb”

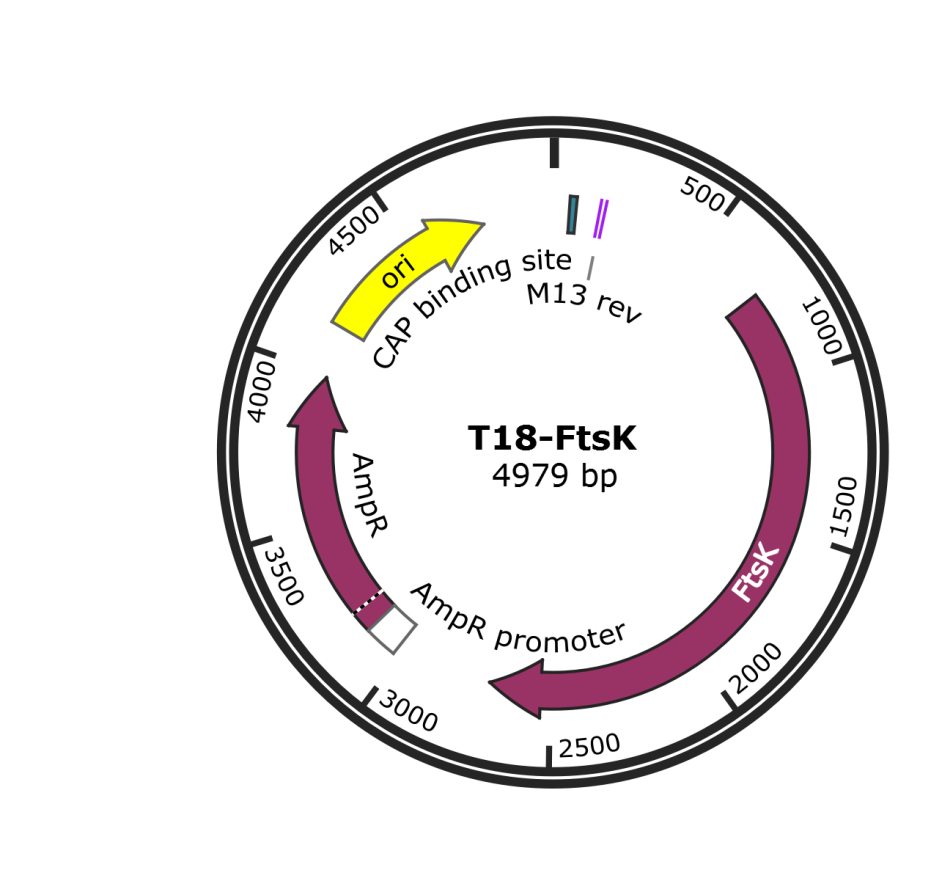

**Figure S18 –** Map of the plasmid T18-FtsK used for analysis of *M. gallisepticum* FtsZ interactions with other proteins. The main elements of the plasmid are shown in the figure. Sequence of the plasmid is shown in file “T18-FtsK.gb”

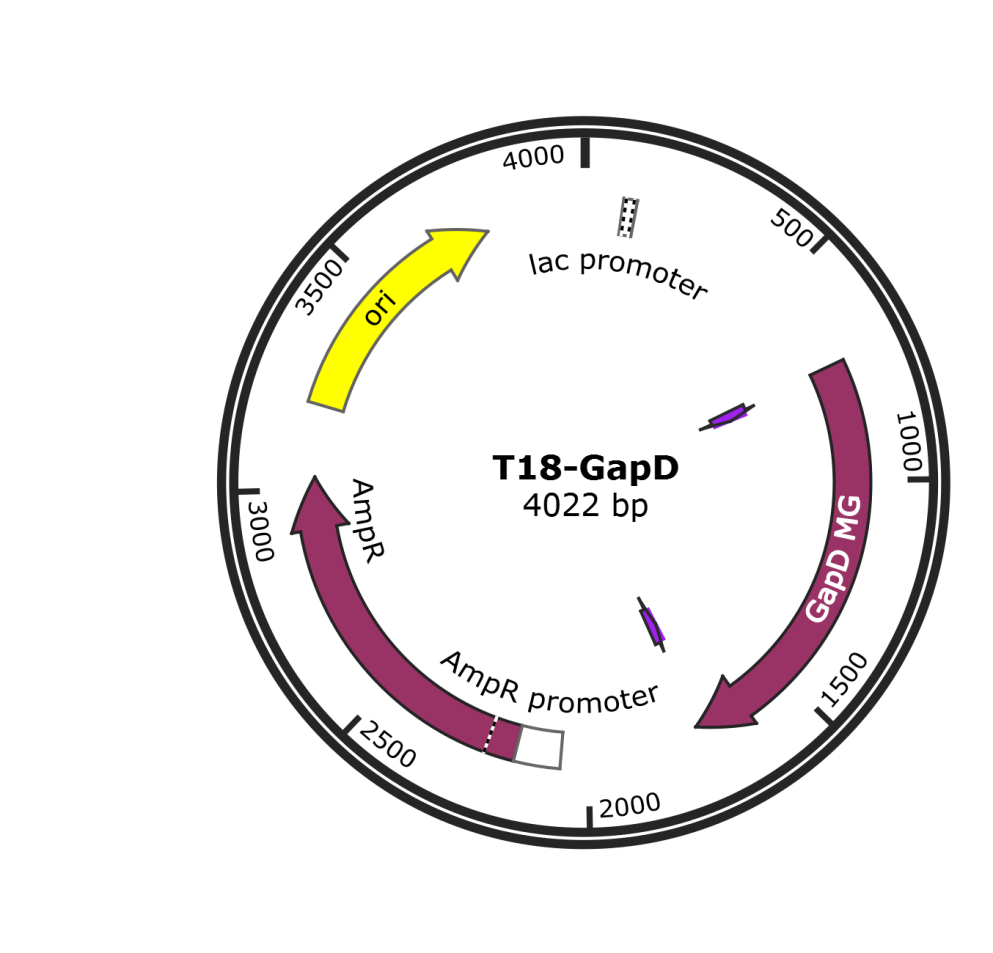

**Figure S19 –** Map of the plasmid T18-GapD used for analysis of *M. gallisepticum* FtsZ interactions with other proteins. The main elements of the plasmid are shown in the figure. Sequence of the plasmid is shown in file “T18-GapD.gb”

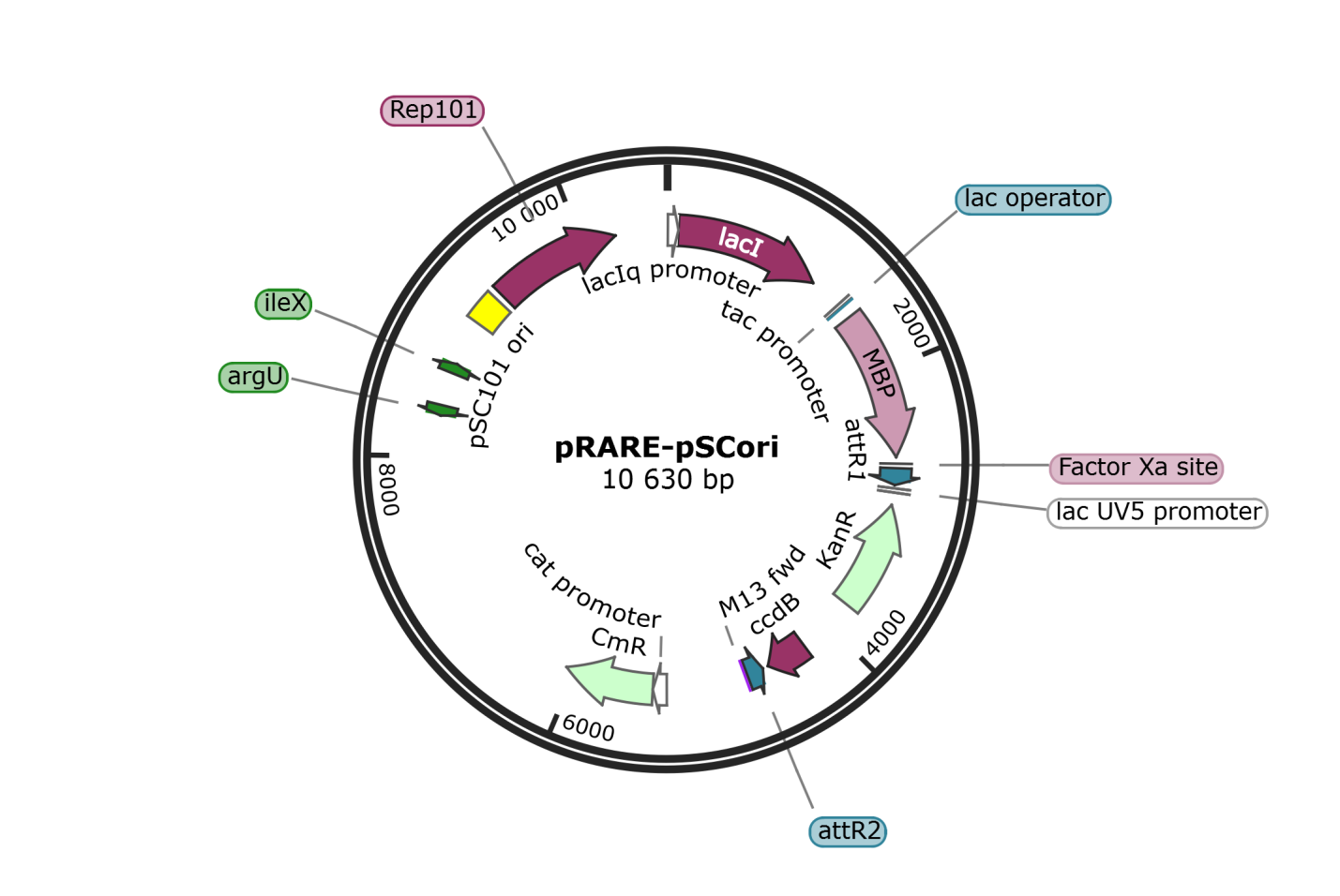

**Figure S20 –** Map of the plasmid pRare-pSCori used to enhance the production of *M. gallisepticum* proteins in *E. coli* for analysis of *M. gallisepticum* FtsZ interactions with other proteins. The main elements of the plasmid are shown in the figure. Sequence of the plasmid is shown in file “pRare-pSCori.gb”

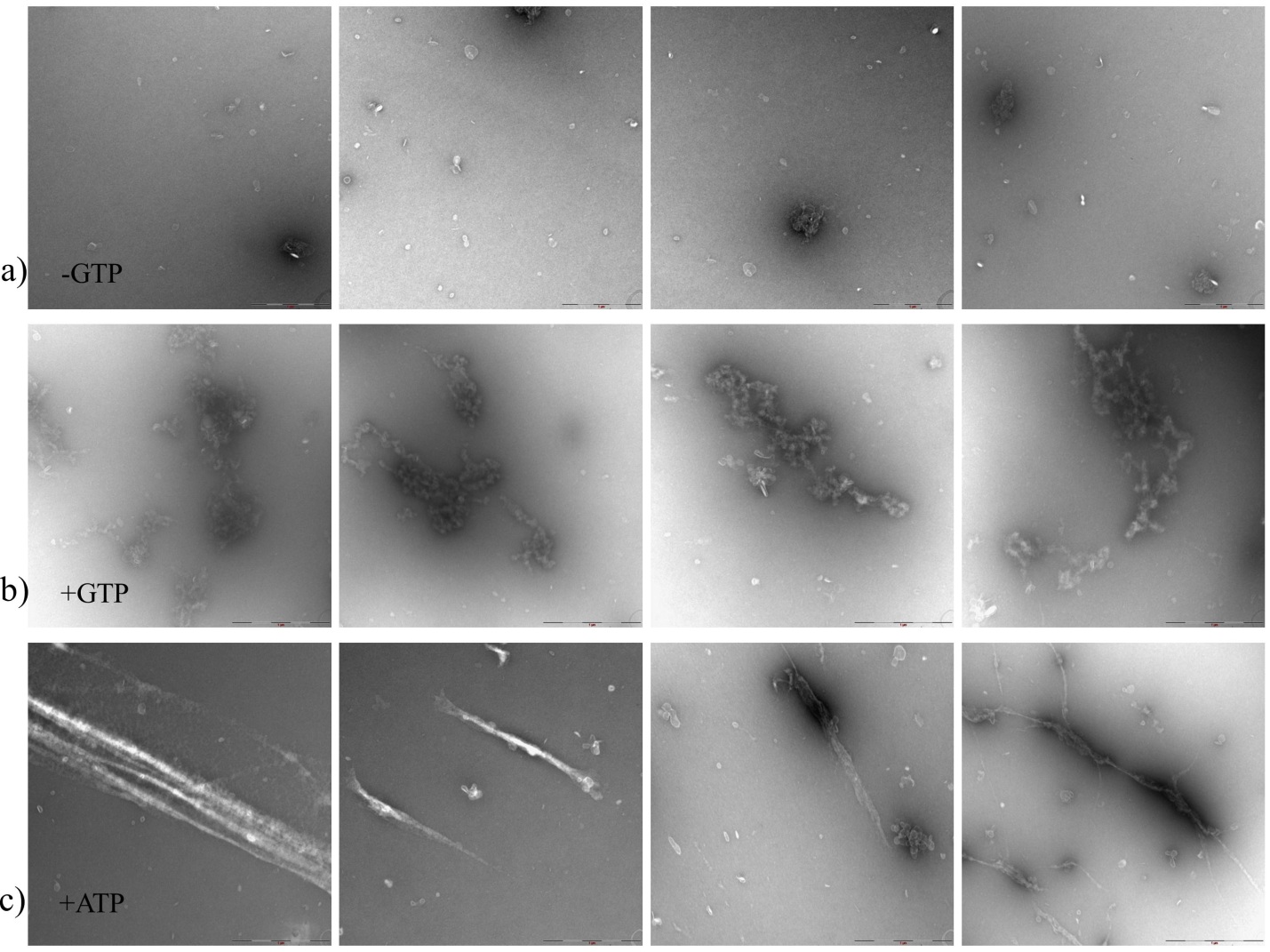

**Figure S21** – *M. gallisepticum* FtsZ polymers visualized by electron microscopy: a) – in the absence of GTP (-GTP); b) – in the presence of GTP (+GTP) and c) – in the presence of ATP

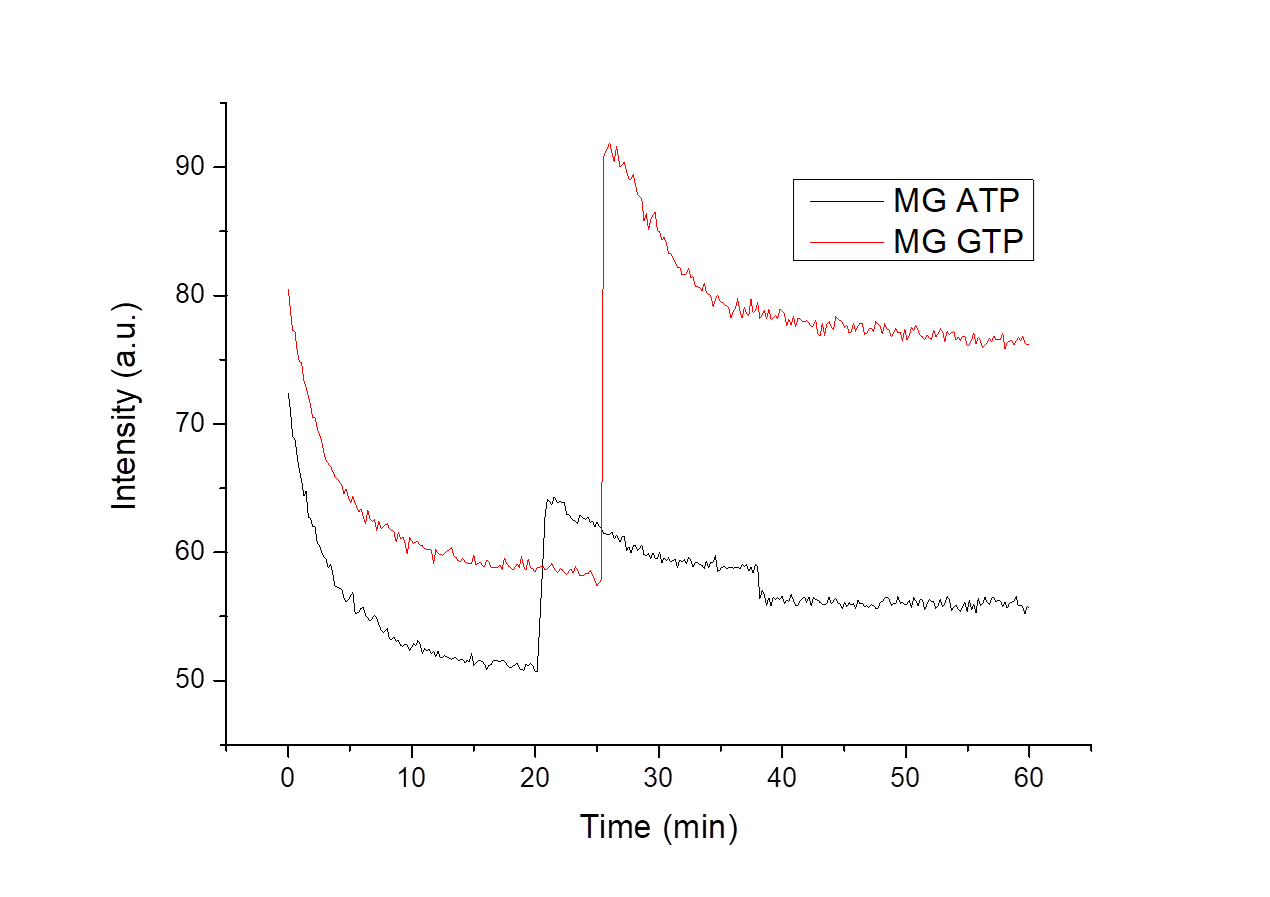

**Figure S22** – Polymerization curves of the *M. gallisepticum* FtsZ protein obtained by static light scattering in the presence of GTP (red line) and ATP (black line). MES-based buffer (pH 6.5) was used. The potassium ion concentration in buffer was 50 mM. The buffers also contained 1 mM EDTA and 5 mM MgCl_2_. The shift in the graphs between 20 and 25 min marks the gap corresponding to the addition of GTP or ATP
